# Temporal mapping of the anti-tumor effects of nanobody-based MSLN.CAR-T cell therapy in metastatic solid tumors

**DOI:** 10.1101/2025.02.26.640438

**Authors:** Chaido Stathopoulou, Mingming Zhao, Qun Jiang, Jessica Hong, Jing Bian, Jingli Zhang, Mitchell Ho, Raffit Hassan

## Abstract

Studies on the dynamic changes occurring in the tumor microenvironment (TME) following CAR-T cell therapy have been confounded by host lymphodepletion, multiple dosing and immunodeficient models. Here, a nanobody-based, mouse mesothelin-targeting CAR-T cell (A101) was developed, achieving effective primary tumor suppression, metastasis reduction, and improved survival after a single dose in immunocompetent, syngeneic mouse models without lymphodepletion. Temporal tumor profiling using RNA sequencing revealed initial downregulation of cell proliferation genes followed by upregulation of inflammation, epithelial-to-mesenchymal-transition (EMT) and extracellular matrix (ECM) modification genes in the CAR-T-treated tumors relative to mock-T-treated controls. This phenotype was reversed at a later timepoint which coincided with downregulation of immunosuppressive *Cd274*^+^ *Lcn2*^+^ neutrophils and upregulation of anti-tumor *P2rx1*^+^ *Nrf2^-^* neutrophils. At the same time, upregulation of *Ccl2*^+^ in fibroblasts and a more immunomodulatory macrophage phenotype was observed in CAR-T-treated tumors, indicating a tumor adaptation mechanism. This study demonstrates complex dynamic changes in the TME, and highlights time-dependent responses of solid tumors to CAR-T cell therapy. It further highlights *Lcn2*^+^ neutrophils and *Ccl2*^+^ fibroblasts as potential therapeutic targets for improving CAR-T cell anti-tumor efficacy.

Graphical abstract.
Tumor responses to A101 CAR-T cell treatment.

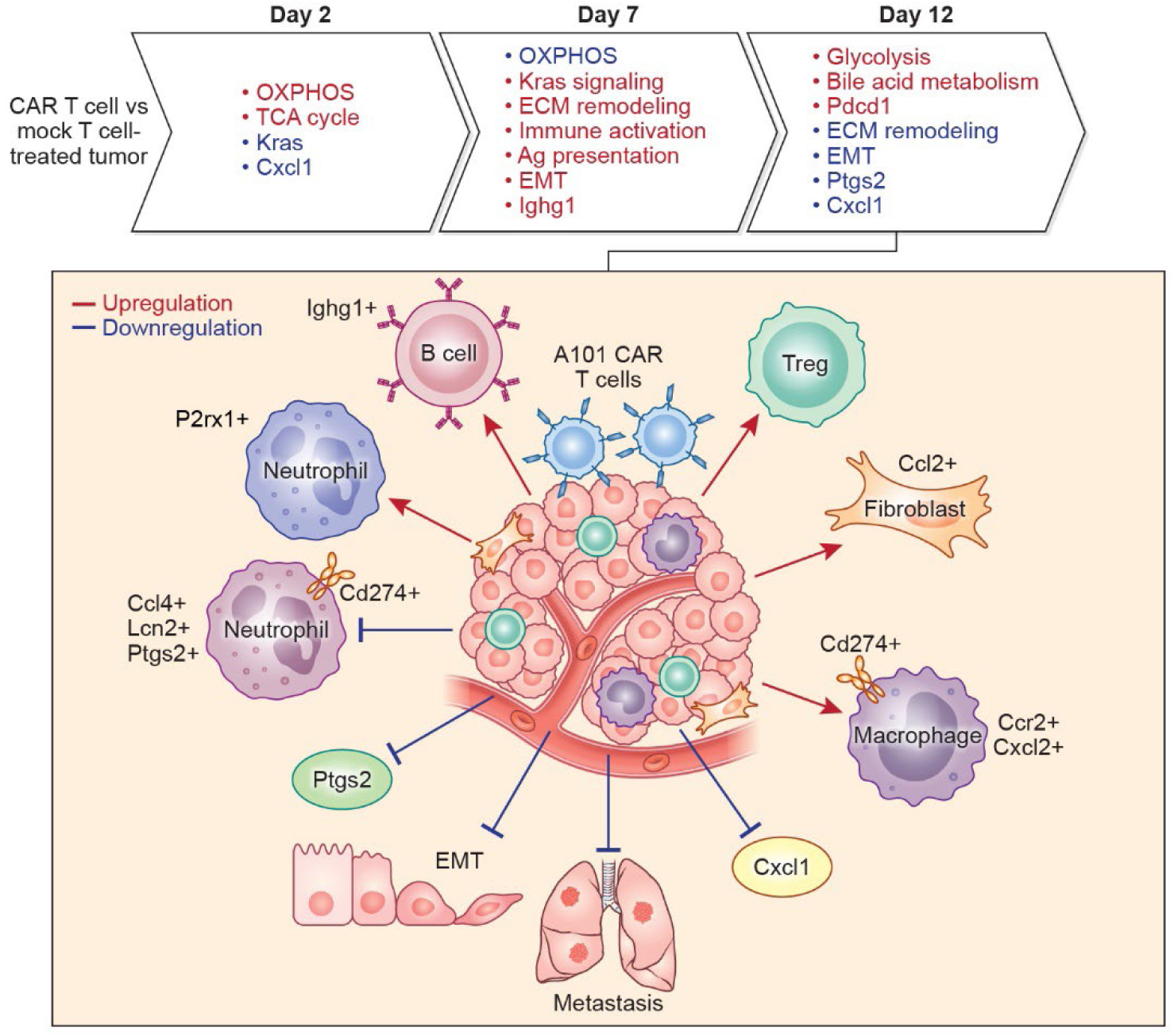

## Introduction

CAR-T cell therapy for solid tumors remains a challenge [1–4]. One major challenge to the development of clinically effective CAR-T cells, is the use of preconditioned hosts and immunodeficient preclinical models which inherently are unsuitable for investigating the role of the endogenous immune system in tumor development, response and resistance to CAR-T cell therapies [5]. The development of mouse CAR-T cells with sufficient anti-tumor efficacy to allow the detailed study of the temporal dynamics of these responses in unprimed hosts is also a challenge. Most mouse-derived CAR-T cells are evaluated in preconditioned hosts [6–8] or models requiring multiple CAR-T cell dosing which can confound mechanistic study results [9, 10]. Another confounding factor can be the use of conventional antibody (Ab) - based CAR structures [6–9, 11]. Conventional Abs are large structures and can present challenges when used in immunocompetent models including increased immunogenicity and protein aggregation due to misfolding [12–14]. Nanobodies on the other hand, offer an attractive alternative owing to their small size, reduced immunogenicity, stability and ability to access epitopes that cannot be accessed by conventional Abs [10, 15, 16].

Here, we sought to develop a mouse nanobody-based CAR-T cell (A101) that can be used in syngeneic models to study temporal changes in the metastatic solid TME following CAR-T cell therapy. The targeted antigen for our A101 CAR-T cells is mouse mesothelin (Msln) which is an antigen expressed in solid tumors such as mesotheliomas, pancreatic, ovarian, and lung carcinomas [17]. It is a malignancy factor that promotes tumor cell proliferation, migration and cell invasion [17]. Its interaction with the CA125/MUC16 ovarian cancer antigen mediates cell adhesion and has been shown to play a role in metastasis of ovarian cancers [17].

To our knowledge, no studies have been performed yet where the entire TME (tumor cells, stroma and immune cells) is analyzed in detail over time, in immunocompetent, unprimed hosts and after only a single dose of mouse CAR-T cells. With our model we were able to use a single dose of CAR-T cells to study TME changes in metastatic solid tumors, in temporal fashion using high throughput techniques including multiparametric flow cytometry, total and single cell RNA sequencing (scRNA-seq). Our study provides new insights into tumor evolution and the dynamic interplay between tumor response and tumor resistance mechanisms and highlights potential targets and timepoints for therapeutic intervention to improve CAR-T cell therapy for solid tumors.

## Results

### A101 nanobody binds to mouse and human mesothelin

Mouse and human mesothelin exhibit a high degree of homology in amino acid sequence and share similarities in both expression patterns and functions (**Fig. 1a**) [18]. Humans and mice typically do not produce antibodies against self-antigens due to negative selection in the thymus. As a result, *in vitro* antibody discovery methods, such as phage display, are used to identify cross-species antibodies targeting human and mouse antigens. Through phage display screening, we identified a camel-derived nanobody A101 which binds to the N-terminal region I of human mesothelin (MSLN) fragment 1AB (**Fig. 1b**). This is the same region that interacts with MUC16 indicating that the nanobodies could potentially block MUC16/mesothelin interaction [19]. Since the amino acid sequence of the mouse and human mesothelin is 71% identical and nanobodies can access small protein regions/epitopes that are otherwise inaccessible to conventional antibodies, we sought to determine whether A101 could also bind to mouse mesothelin [15, 18]. Using flow cytometry, specific binding of A101 to mouse and human mesothelin was confirmed on several mouse and human-derived tumor cell lines (**Fig. 1c**). Low binding was observed on the mouse-derived lung adenocarcinoma cell line TC-1 (Msln^low^). No binding was observed on the negative control cell lines which included the human-derived Lewis lung carcinoma cell line (LLC) and the pancreatic carcinoma cell line KLM1 MSLN knockout (KO) (**Fig. 1c**).

**Fig. 1.**
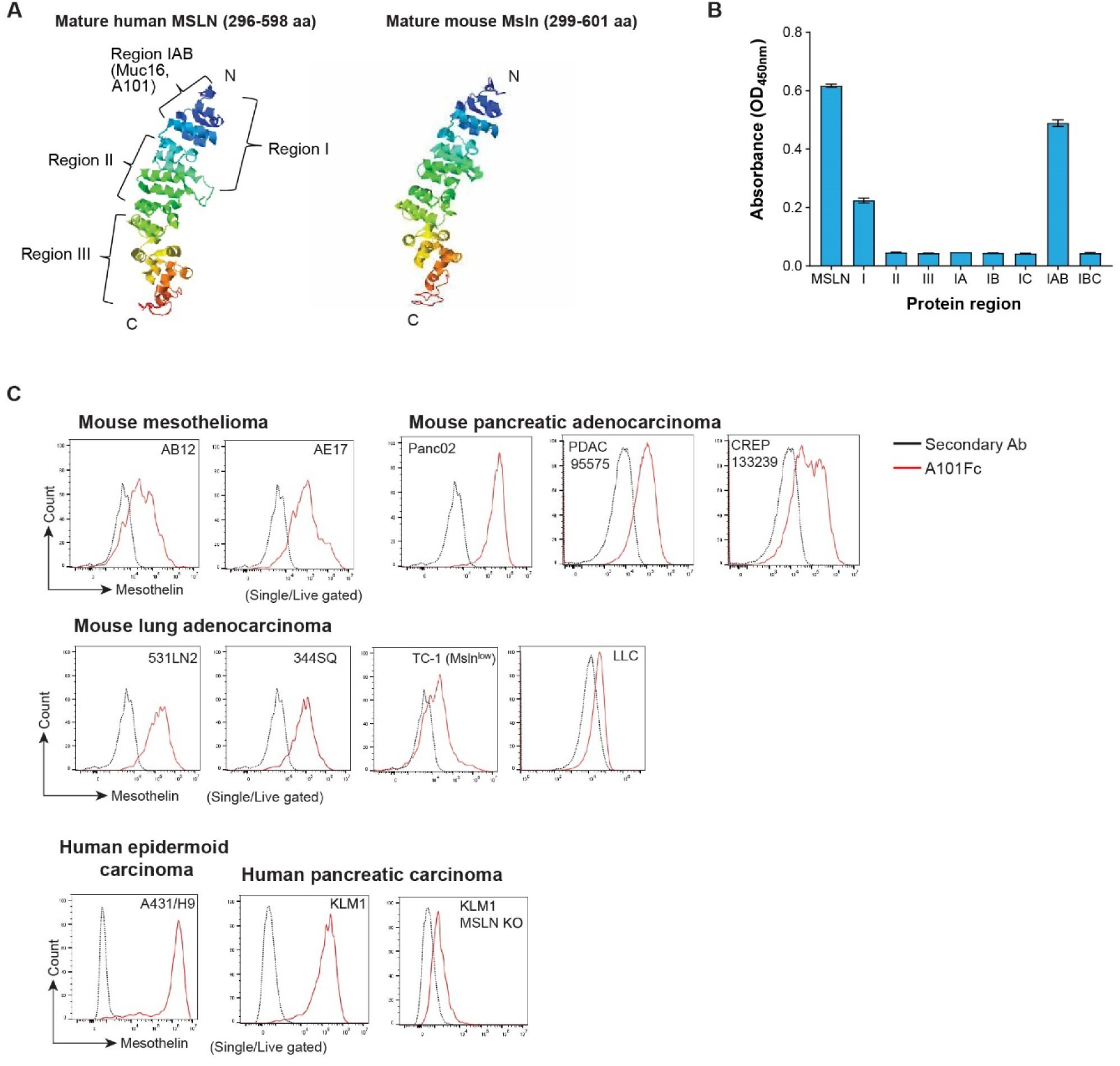
Epitope binding of A101 V_H_H on mesothelin. **a** Protein structure of mature human and mouse mesothelin. The protein structure model was built by I-TASSER software based on the mesothelin sequence. **b** ELISA assay measuring binding of A101 V_H_H to human mesothelin fragments and full length mesothelin (MSLN). **c** Histogram of mesothelin expression on murine- and human-derived tumor cell lines analyzed by flow cytometry using A101Fc and anti-human secondary antibody.

Collectively, these experiments showed that A101 can bind to both mouse and human mesothelin while maintaining antigen specificity indicating it is a suitable candidate for the development of mouse Msln.CAR-T cells.

### Mouse A101 CAR-T cells show anti-tumor activity and antigen specificity *in vitro*

We generated a second-generation CAR vector consisting of the A101 V_H_H linked to a murine CD8 transmembrane/hinge domain and murine 4-1BB and CD3ζ signaling domains along with a truncated human EGFR protein (EGFRt) as a surface marker for tracking CAR expression (**Fig. 2a**). The CAR codon sequence was optimized for the mouse host. To generate CAR-T cells, CD3^+^ T cells were isolated from mouse splenocytes, activated, transduced and expanded *in vitro*. Cell cultures were optimized, and we were able to achieve greater than 60% transduction efficiency (**Fig. 2b, Supplementary Fig. 1a, b**). Transduced and mock-T cells showed similar proliferation levels following stimulation with αCD3/αCD28 beads as well as similar levels of exhaustion markers at the end of the culture (**Fig. 2c, d**). The final products were also similar in T cell subset composition (**Supplementary Fig. 1c, d**). Collectively, this indicates that the cell processing following lentiviral transduction had no significant impact on the cell phenotype and proliferative capacity of the CAR-T cells.

**Fig. 2.**
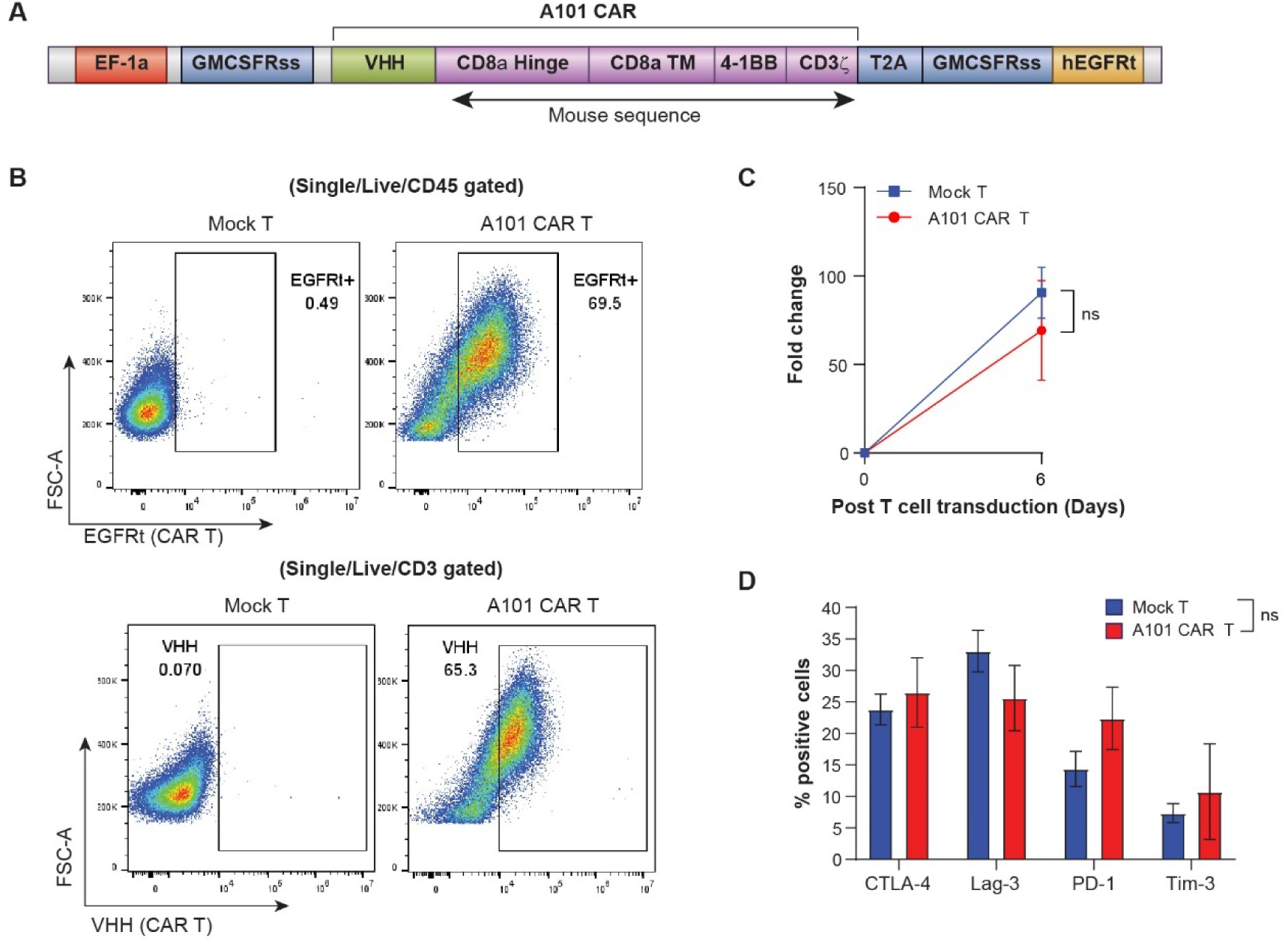
Proliferation and phenotypic characterization of A101 CAR-T cells *in vitro*. **a** Depiction of A101 CAR construct. **b** Transduction efficiency of mouse A101 CAR-T cells measured by flow cytometry. CAR-binding PE-conjugated anti-human EGFR antibody (top panels) or APC-conjugated anti-camel V_H_H antibody was used. Gating strategy is shown in **Supplementary Fig. 1a. c** Comparable proliferation of A101 CAR-T and mock-T cells induced by anti-CD3/28 beads *in vitro* (n = 5 independent samples/group). Data indicate mean ± SE and groups were compared using two-side Student’s t test (t=0.6787, df=8). No significant difference was found. **d** Characterization by flow cytometry of A101 CAR-T and mock-T cell expression of CTLA-4, Lag-3, PD-1 and Tim-3 on day 6 post-transduction (n = 6 independent samples/group). Cells were gated on CD3^+^ cells. Data indicate mean ± SE and groups were compared using two-way ANOVA with Sidak’s multiple comparisons test (F=0.4168 (CTLA-4), 1.162 (Lag-3), 1.246 (PD-1), 0.5288 (Tim-3), df=40)). No significant differences were found (p>0.05). Gating strategy for (**d**) is shown in **Supplementary Fig. 1d**.

To determine whether the transduced cells maintain their effector function and antigen specificity, we assessed their cytolytic capacity in coculture assays with tumor cells expressing mesothelin. A101 CAR-T cells were cocultured with the murine-derived cell lines 344SQ, AE17, and 531LN2 as well as the negative control cell line TC-1 (**Fig. 3a-d**). Antigen specificity was assessed by comparing A101 CAR-T cell effector function to that of mock-T cells. Contrary to mock-T cells, A101 CAR-T cells consistently exhibited dose-dependent and antigen-specific cytotoxicity with ∼50% killing observed at effector to target (E:T) ratio of 25 or higher (**Fig. 3a-d**). The mouse-derived pancreatic cell lines PDAC 95575 and CREP133239 which are Msln positive, were also shown to be sensitive to A101 CAR-T cell-mediated killing in a dose-dependent manner (**Fig. 3e, f, Supplementary Fig. 2a-d)**. Expression levels of the effector cytokines IL-2, IFN-γ and TNF-α were assessed 24 hours after coculture. A101 CAR-T cells expressed higher levels of these effector cytokines compared to mock-T cells (**Fig. 3g, h, Supplementary Fig. 2e-g**). Interestingly, a lower E:T ratio was required for 50% killing when human A101 CAR-T cells were cocultured with human tumor cell lines (**Supplementary Fig. 2h-k**). In fact, their killing capacity was comparable to that of human YP218 CAR-T cells *in vitro* (**Supplementary Fig. 2h-k**) [20], which indicates that A101 is capable of strong binding to its epitope. This combined with the flow data, indicates that the affinity level of A101 binding to mouse Msln does not play a role in the higher E:T ratios required in the mouse coculture experiments.

**Fig. 3.**
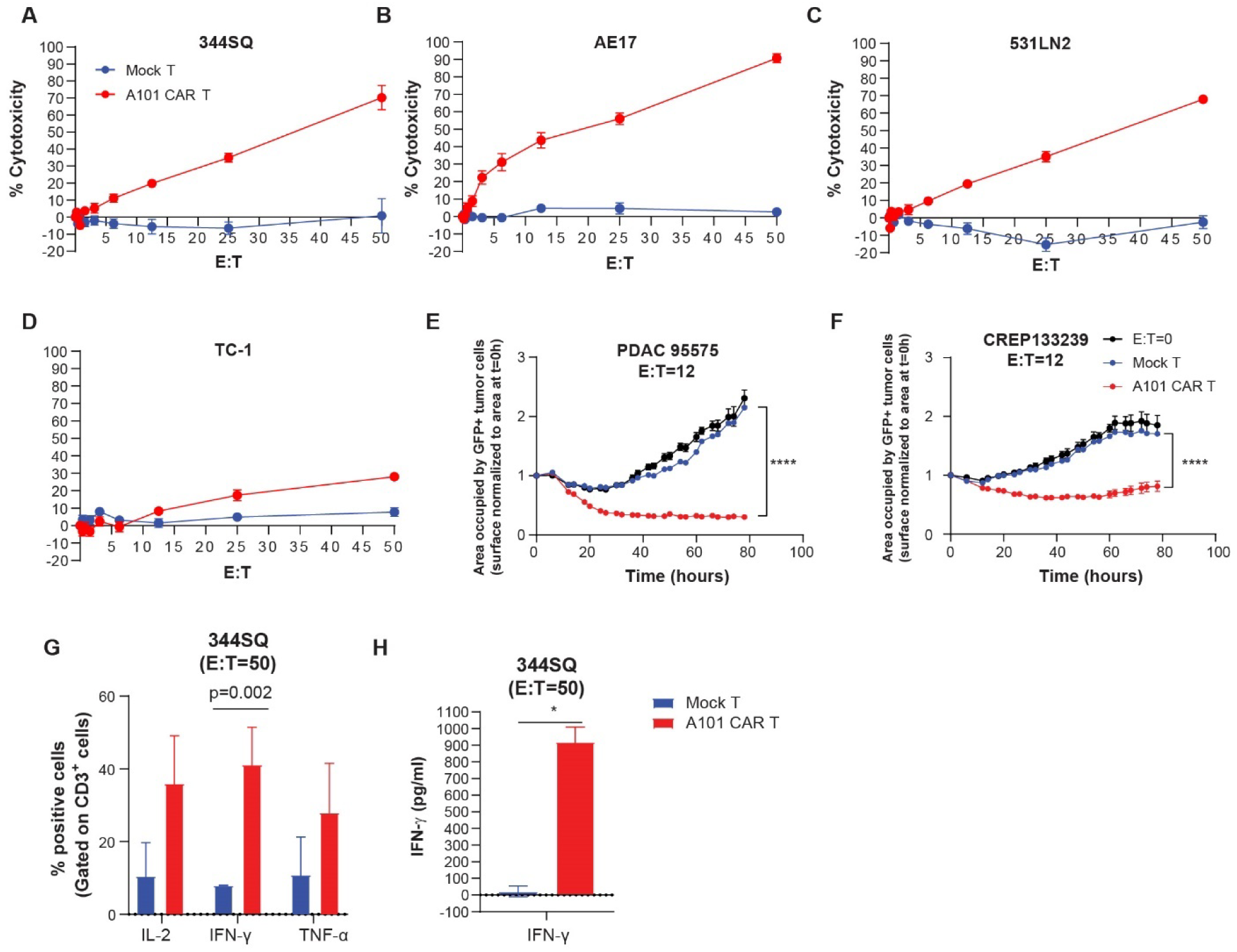
Functionality of A101 CAR-T cells *in vitro*. **a-d** Cytotoxic activity of T cells when co-cultured with tumor cells expressing the targeted antigen. The cell lines tested were 344SQ (**a**), AE17 (**b**), 531LN2 (**c**) tumor cells and Msln^-/low^ TC-1 cells (**d**). n=8 independent experiments except for TC-1 (n=4). Mean ± SE is shown. Statistical analysis was performed using unpaired, two-side Student’s t test. For A101 CAR-T vs mock-T; *p=0.0351 (344SQ, t=2.279, df=18), *p=0.0203 (AE17, t=2.546, df=18), *p=0.0350 (531LN2, t=2.280, df=18). No other significant differences were found. **e-f** Cytotoxic activity of T cells during 78hrs of co-culture with GFP^+^ Msln^+^ murine tumor cell lines PDAC 95575 (**e**) and CREP133239 (**f**) at E:T ratio of 12 (n=3). Mean ± SE is shown. Groups were analyzed using two-way ANOVA with Tukey’s multiple comparisons test (for PDAC 95575: F=464.7, df=6, for CREP133239: F=119.9, df=6). ****p<0.0001. **g** Flow cytometric quantification of A101 CAR-T and mock-T cell expression of IFN-γ, IL-2 and TNF-α at E:T=50, 24hrs after coculture with 344SQ. n = 2. Mean ± SE is shown. Statistical analysis was performed using unpaired, two-side Student’s t test (t=1.599 (IL-2), 3.253 (IFN-γ), 1.003 (TNF-α), df=2)). No significant difference was found. Gating strategy is shown in **Supplementary Fig. 2e. h** Quantification of IFN-γ secretion in co-culture supernatant at E:T=50, 24hrs after coculture with 344SQ cells using ELISA. n = 2. Mean ± SE is shown. Statistical analysis was performed using unpaired, two-side Student’s t test. *p=0.0111 (t=9.387, df=2).

Collectively, this data demonstrates that mouse T cells can be effectively transduced using a lentiviral vector and the A101 CAR-T cells are able to expand *in vitro* while maintaining their effector function and antigen-specificity.

### Mouse A101 CAR-T cells show superior anti-tumor efficacy in syngeneic lung and pancreatic adenocarcinoma models

For our preclinical models, we utilized syngeneic mouse models of metastatic lung adenocarcinoma (344SQ) and pancreatic cancer (PDAC 95575). Both cell lines are Msln positive and derive from KRAS mutant mouse strains; 344SQ from *p53R172^HΔg/+^K-ras^LA1/+^*mice and PDAC 95575 from *LSL-Kras^G12D/+^; LSL-Trp53^R172H/+^; Pdx-1-Cre* (KPC) mice.

To assess the *in vivo* effector function of A101 CAR-T cells, we treated 344SQ tumor bearing mice with 5 x 10^6^ CAR-T cells and an equal total number of mock-T cells or saline intravenously. Compared to saline or mock-T cells, A101 CAR-T cell treatment significantly reduced primary tumor growth rate (**Fig. 4a**) and improved survival rates (**Fig. 4b**). Median survival was 36.5 days for the CAR-T-treated group, 20 days for the mock-T and 27 days for the saline group (**Fig. 4b**). More importantly, reduced metastases were observed in the lungs of the CAR-T-treated group where <1% of the lung tissue had metastasis by day 33 after treatment (**Fig. 4c, d**). Collectively, this indicates that despite the high E:T ratios required for sufficient killing i*n vitro*, a single dose of A101 CAR-T cells is sufficient to significantly reduce primary tumor growth rate, prevent or delay lung metastasis and improve survival. In addition, no toxicity was observed in the treated mice and normal body weight was maintained throughout treatment. This indicates that A101 CAR-T cells do not bind to normal mouse tissues that express mesothelin and therefore are non-toxic.

**Fig. 4.**
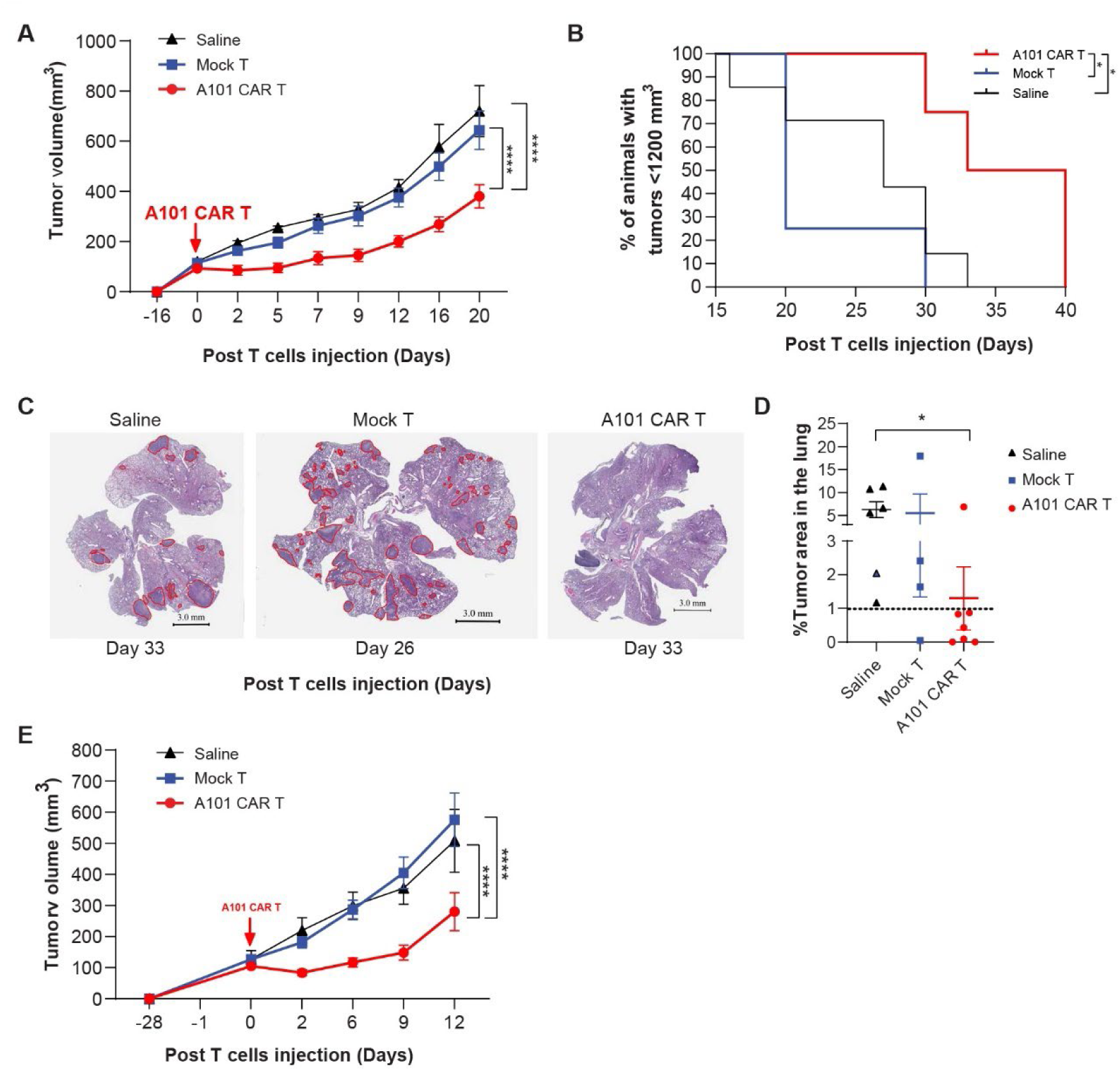
Functionality of A101 CAR-T cell *in vivo*. **a** Syngeneic 344SQ tumor-bearing 129sv mice were treated with one dose of 5 x 10^6^ A101 CAR-T or mock-T cells or saline. Average tumor growth for all cohorts. Data indicate mean ± SE and groups were compared using two-way ANOVA analysis with Tukey’s multiple comparison tests (n = 6 for A101 CAR-T, n = 7 for mock-T and saline). ****p<0.0001 (F=41.42, df=2). **b** Kaplan-Meier curves of 344SQ bearing mice from each of the indicated treatment groups. Data points are mean ± SE (n = 4 for the A101 CAR-T and mock-T, n = 7 for saline) and groups were compared using the log-rank test. For A101 CAR-T vs. mock-T; *p=0.0171, A101 CAR-T vs. saline; *p=0.0253. Median survival in days for each group is: 36.5 (CAR-T), 20 (mock-T), 27 (saline). **c** In a separate experiment, syngeneic 344SQ tumor-bearing 129sv mice were treated with one dose of 3 x 10^6^ A101 CAR-T or mock-T cells or saline. Lungs were harvested 3-6 weeks (Day 26 – 40) post T cells injection. Representative images showing the presence of metastatic lesions in the lung outlined in red on day 33 (A101 CAR-T and saline) and day 26 (mock-T) post T cells injection. Scale bar: 3 mm. **d** Percent lung area that is metastatic for all cohorts. Data points are mean ± SE (n = 7 for A101 CAR-T, n = 4 for mock-T, n = 6 for saline). Groups were compared using two-side Student’s t test. *p=0.0237 (t=2.622, df=11). No other significant differences were found (p>0.05). **e** Treatment of PDAC 95575 syngeneic tumor bearing C57BL/6NCrL mice with 4 x 10^6^ A101 CAR-T cells, mock-T cells or saline. Average tumor growth for all cohorts. Data indicate mean ± SE and groups were compared using two-way ANOVA analysis with Tukey’s multiple comparison tests (n = 10 for A101 CAR-T and mock-T, n = 7 for saline). ****p<0.0001 (F=23.23, df=2).

To validate our findings, we sought to determine whether A101 CAR-T cells can control tumor growth in a second tumor model. The pancreatic cell line PDAC 95575 showed increased mesothelin expression and sensitivity to A101 CAR-T cell mediated killing *in vitro* (**Fig. 3e**). Therefore, we established syngeneic tumors in *C57BL/6* mice and treated them with 4 x 10^6^ CAR-T cells and an equal total number of mock-T cells or saline (**Fig. 4e**). A101 CAR-T cells were able to significantly reduce PDAC 95575 tumor growth rate (**Fig. 4e**). However, due to the aggressiveness of the model, rapid tumor ulceration necessitated the termination of experiments at an early timepoint (day 12-15 post-treatment).

In summary, a single dose of A101 CAR-T cells was able to suppress tumor growth without lymphodepletion in two different tumor models.

### A101 CAR-T cells home to the tumor within 48hrs post-injection leading to an increase of CD8 T cells, B cells and Treg cells in the TME

Since A101 CAR-T cells were able to inhibit tumor growth after a single dose, we hypothesized that A101 CAR-T cells can traffic to the tumor and alter the TME. To study these changes, we analyzed tumor tissue and blood using flow cytometry at different timepoints after treatment (**Fig. 5a-d, Supplemental Fig. 3a-c**).

**Fig. 5.**
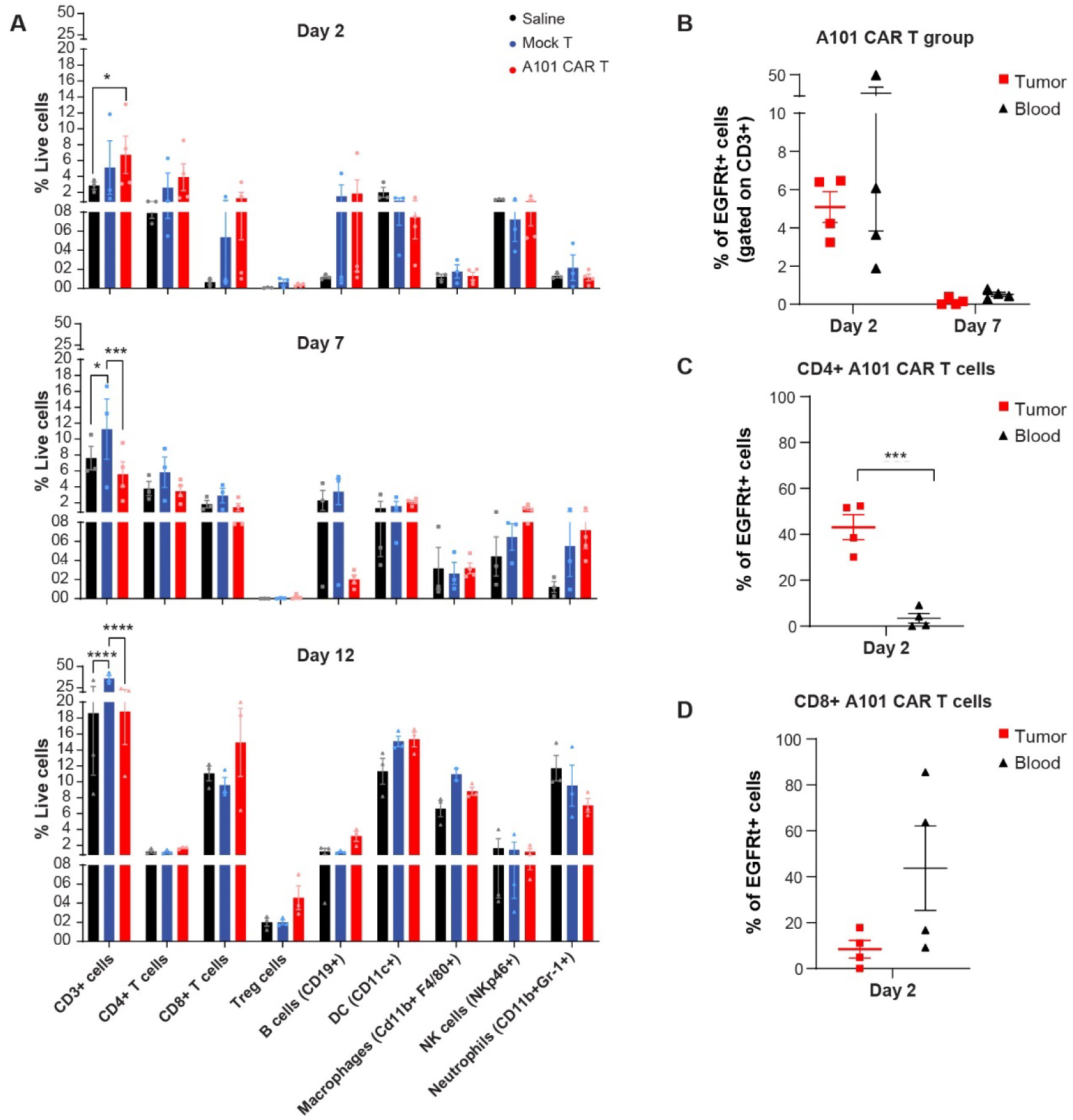
Characterization of TME changes in response to A101 CAR-T cell treatment. **a-e** Syngeneic 344SQ tumor-bearing 129sv mice were treated with one dose of 3-5 x 10^6^ A101 CAR-T or mock-T cells or saline. Tumors were harvested on day 2, 7 and 12 post T cells injection. Blood was harvested on day 2 and 7. Live cells from tumor tissue were column sorted prior to flow cytometry. **a** Cell population analysis of tumors harvested on day 2, 7 and 12 post T cells injection. Data points are mean ± SE (n = 4 for day 2-7 A101 CAR-T, n=2 for day 12 mock-T macrophages, n = 3 for the rest) and groups were compared using two-way ANOVA analysis with Tukey’s multiple comparison tests. For day2; *p=0.0351 (F=1.725, df=2), for day 7; ***p=0.0003, *p=0.0416 (F=3.912, df=2), for day 12; ****p<0.0001 (F=2.587, df=2). **b** The percentage of EGFRt^+^ CAR-T cells among CD3^+^ cells in A101 CAR-T-treated tumors and blood. **c-d** The percentage of CD4^+^ (**c**) and CD8^+^ (**d**) EGFRt^+^ CAR-T cells in tumors and blood of the A101 CAR-T-treated group is shown. Data points are mean ± SE (n = 4 for the A101 CAR-T group, n = 3 for controls) and groups were compared using unpaired, two-side Student’s t test. ***p=0.0005 (t=6.872, df=6). Gating strategy for (**a**) is shown in **Supplementary Fig. 3b**. Gating strategy for (**b-d**) is shown in **Supplementary Fig. 3a**.

For 344SQ tumor characterization, we harvested tissues on day 2, 7 and 12 post-treatment and analyzed the immune cell populations (**Fig. 5a, Supplemental Fig. 3a, b**). Over time, in all the treatment groups, there was an increase in myeloid cell populations (macrophages, dendritic cells (DCs), neutrophils) and lymphocytes (CD3 T cells, CD8 T cells and Treg cells) (**Fig. 5a**). No changes in total CD4 T cells or NK cells were observed (**Fig. 5a**). However, when compared to mock-T, A101 CAR-T-treated tumors exhibited decreased levels of CD3 T and B cells and an increase in NK cells on day 7 post-treatment (**Fig. 5a**). On day 12, they showed an increase in CD8 T, B cells and Treg cells but a decrease in total CD3 T cells and macrophages (**Fig. 5a**).

We also assessed CAR-T cell trafficking, persistence and phenotype in tumors and blood isolated on days 2 and 7 post-treatment (**Fig. 5b-d, Supplemental Fig. 3a, c**). Peak levels of A101 CAR-T cells were detected in tumors and blood on day 2 post-treatment with levels rapidly reduced by day 7 (**Fig. 5b, Supplemental Fig. 3a, c**). Interestingly, most A101 CAR-T cells in the 344SQ tumors on day 2 were CD4^+^ T cells whereas in the blood, most of them were CD8^+^ (**Fig. 5c, d, Supplementary Fig. 3a**).

In the PDAC 95575 model, tumors were harvested on day 2 and 15 post-treatment and characterized by flow cytometry. A similar pattern was observed where peak CAR-T cell levels were observed on day 2 post-treatment (**Supplemental Fig. 3d**). Over time, in all groups, there was a decrease in macrophages, CD4 T, CD8 T and Treg cells (**Supplemental Fig. 3e**). Compared to mock-T, CAR-T cell-treated tumors showed a decrease in B, DC and NK cells on day 2 (**Supplemental Fig. 3e**). However, on day 15, there was an increase in DC, macrophages and CD8 T cells (**Supplemental Fig. 3e**).

These results indicate that A101 CAR-T cells can traffic to the tumor within 48hrs post-inoculation and despite their low persistence, they can induce changes in the TME potentially altering tumor growth and metastasis.

### TME characterization using total RNA-seq shows increase of plasma and Treg cells in A101 CAR-T-treated tumors

To analyze the tumor immune landscape in more detail, we performed total RNA-seq of tumors isolated on day 2, 7 and 12 post-treatment and performed CIBERSORT analysis using the deconvoluted LM22 signature matrix (**Fig. 6a-c**) [21]. In this experiment mice were treated with one dose of 8 x 10^6^ A101 CAR-T cells or an equal number of total mock-T cells (**Fig.6a**). Transcriptomic analysis showed that over time, in all the treatment groups, there was an increase in M2 macrophages, memory B cells, plasma cells, Tfh cells and Treg cells (**Fig. 6b**). No significant changes were observed in total CD4 T, NK, CD8 T, naïve B cells, neutrophils, eosinophils, monocytes, M0 and M1 macrophages, DCs or mast cells (**Fig. 6b, c**).

**Fig. 6.**
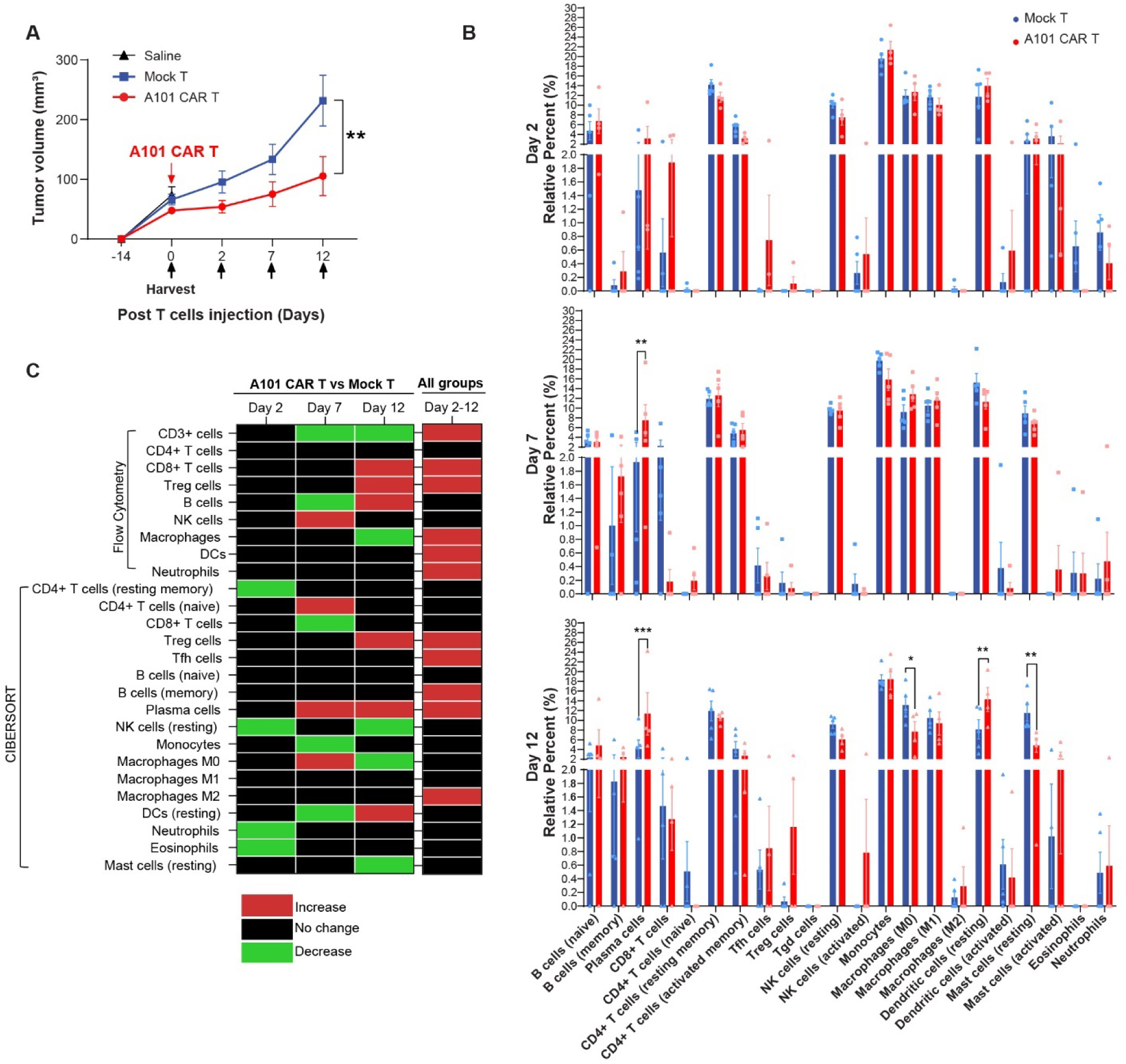
Tumor characterization using total RNA-seq. **a** Syngeneic 344SQ tumor-bearing 129sv mice were treated with one dose of 8 x 10^6^ A101 CAR-T or mock-T cells or saline. Tumors were harvested on day 0 for untreated control (saline) or on day 2, 7 and 12 post-administration of T cells for the A101 CAR-T and mock-T groups and transcriptomic analysis was performed. Average tumor growth for all cohorts. Data indicate mean ± SE (n = 4 for A101 CAR-T, n = 5 for mock-T, n = 5 Saline). Groups were compared using two-way ANOVA with Sidak’s multiple comparisons test (F=6.341, df=2). **p=0.0027. **b** CIBERSORT cell abundance analysis using the LM22 deconvoluted signature matrix. Data points are mean ± SE and groups were compared using two-way ANOVA analysis with Tukey’s multiple comparison tests (n = 4 for A101 CAR-T on day 2 and day 12, n = 5 for the rest of the groups). F=7.749e-015, df=5. For Day 7; **p=0.0071, ***p=0.0003, *p=0.0154, Day 12 dendritic cells resting A101 CAR-T vs mock-T; **p=0.0044, Day 12 mast cells resting A101 CAR-T vs mock-T; **p=0.0014. **c** Summary of dynamic changes in immune cell populations observed in the A101 CAR-T-treated group relative to mock-T control using flow cytometry and CIBERSORT analysis. Cell population changes observed over time (day 2 to 12) in all groups regardless of treatment are also shown (the flow cytometry data include: saline, mock-T, CAR-T groups, the CIBERSORT data include: mock-T and CAR-T groups). Black color indicates no change, red indicates increase, green indicates decrease.

Compared to mock-T control, on day 2 post-treatment, CAR-T cell-treated tumors showed decreased levels of resting memory CD4 T cells, resting NK cells, eosinophils and neutrophils (**Fig. 6b, c**). On day 7, they showed increased levels of plasma cells, naive CD4 T cells and M0 macrophages (**Fig. 6b, c**). CD8 T cells, monocytes and resting DCs were decreased (**Fig. 6b, c**). On day 12, CAR-T cell-treated tumors showed increased levels of plasma cells, Treg cells and resting DCs compared to mock-T control (**Fig. 6b, c**). Resting mast cells, M0 macrophages and resting NK cells were decreased (**Fig. 6b, c**).

No significant differences were observed in total CD4 T cells or NK cells which was consistent with the flow data (**Fig. 6c**). Treg cells in the CAR-T group increased over time relative to control in both the flow cytometry and CIBERSORT analysis (**Fig. 6c**). Another similarity found, was the increase in B cells (flow) and plasma cells (CIBERSORT) in the CAR-T group relative to mock-T on day 12 post-treatment (**Fig. 6c**). Signaling pathway analysis also showed upregulation of B cell activation in the CAR-T cell-treated tumors (**Supplemental Fig. 4a**).

### Tumors upregulate EMT and ECM remodeling pathways in a time dependent manner

To better understand the temporal changes in the TME following CAR-T cell treatment, we performed signaling pathway analysis from the same transcriptomic data. Metabolic changes were observed in the CAR-T-treated tumors over time that were distinct form the mock-T control.

Specifically, two days after treatment, CAR-T cell-treated tumors upregulated fatty acid and galactose metabolism, OXPHOS and TCA cycle pathways (**Fig. 7a, b**). In addition, they upregulated amino acid degradation pathways (e.g. valine, leucine and lysine degradation pathways) (**Fig. 7a**) and downregulated cell cycle and cell proliferation related pathways as well as *Mki67* and *Kras* expression compared to mock-T-treated tumors (**Fig. 7a, b**). This result suggests that CAR-T treatment successfully disrupted tumor proliferation in its early stages, forcing the tumor to undergo metabolic reprogramming to meet the energy and biosynthetic demands required for survival under immune-mediated stress. This was consistent with the reduced primary tumor growth rate observed *in vivo*.

**Fig. 7.**
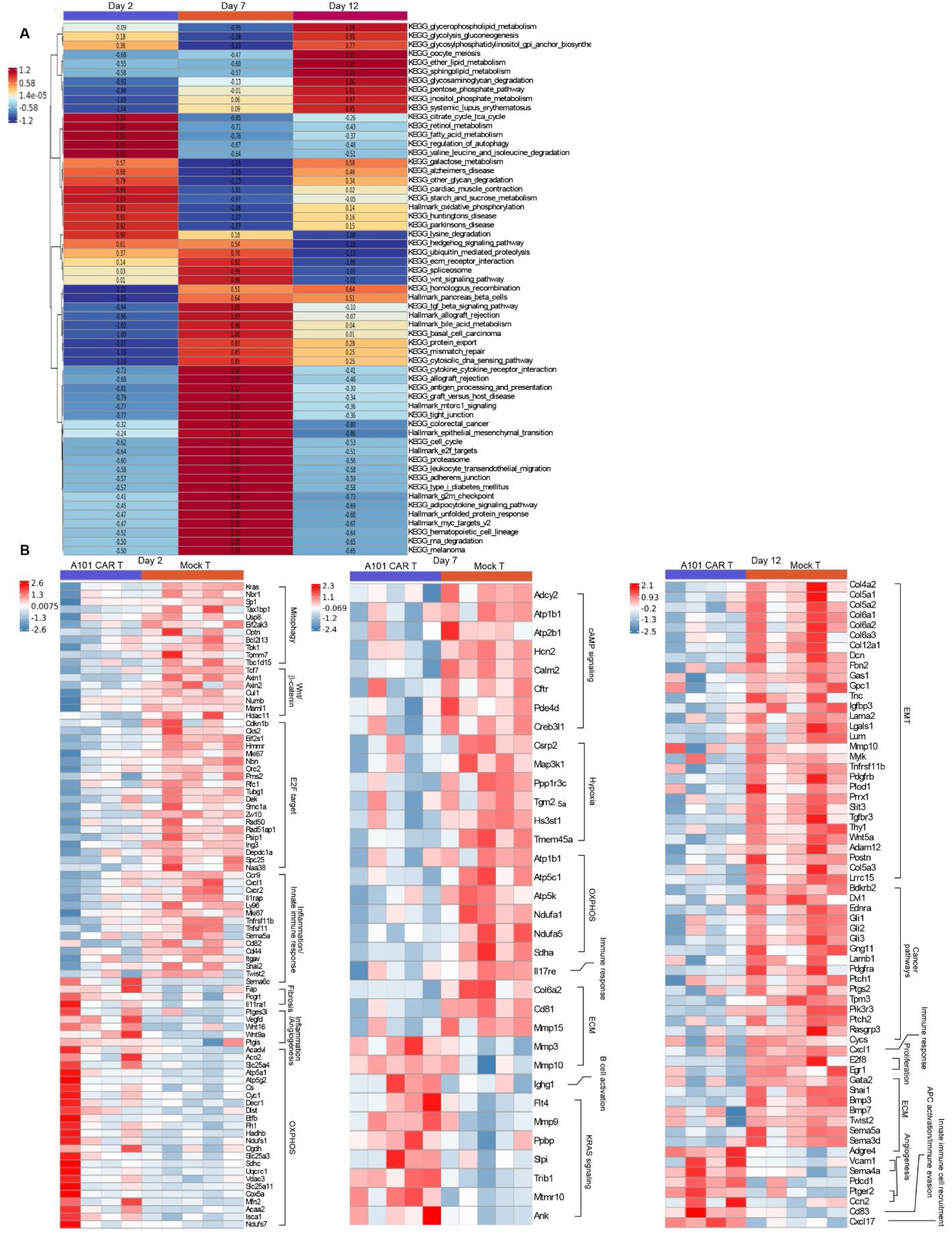
Signaling pathway analysis using total RNA-seq. **a** Heatmap of difference in pathway activity in A101 CAR-T tumors relative to mock-T control on day 2, 7 and 12 post T cells injection. GSVA scores were averaged for n = 4-5 donors/group (n = 4 for A101 CAR-T on day 2 and day 12, n = 5 for A101 CAR-T on day 7, n = 5 for mock-T on day 2, 7 and 12). Enrichment scores of top pathways for the A101 CAR-T groups were calculated as ratios relative to the scores of mock-T and are reported in each cell. **b** Heat map of significant DEGs (pval<0.05) associated with signaling pathways identified from the L2P Fisher’s Exact Test analysis.

In addition to the metabolic changes, genes associated with inflammation such as *Vegfd, Ptges3l* and *Ptgis* were also found upregulated at this timepoint (**Fig. 7b**). However, immune response genes such as *Cxcl1, Ccr9, Cxcr2, Cd44, Cd82, Ly96* and *Tcf7* were significantly downregulated in CAR-T-treated tumors (**Fig. 7b**).

On day 7 post-treatment, major ATP-synthesis pathways associated with cell proliferation were significantly downregulated in the CAR-T-treated tumors compared to mock-T controls. These included the pentose-phosphate, TCA cycle, OXPHOS and glycolysis-glucogenesis pathway (**Fig. 7a**). This change was accompanied by significant upregulation of ECM modification pathways such as the adherens junction, tight junction, EMT, TGF-β signaling, ECM receptor interaction and protein export pathway (**Fig. 7a**). ECM modification correlated with significant upregulation of inflammatory and immune activation pathways. These included immune cell recruitment pathways (leucocyte transendothelial migration), innate immune response pathways (cytosolic DNA sensing), antigen presentation-associated pathways (antigen processing and presentation, allograft rejection, graft versus host disease) and other immune response pathways such as the hematopoietic cell lineage, adipocytokine signaling and cytokine-cytokine receptor interaction pathway (**Fig. 7a**). Consistent with this observation is the upregulation of the B cell activation gene *Ighg1* (**Fig. 7b**).

On day 12 post-treatment, CAR-T-treated tumors significantly upregulated glucose metabolism pathways compared to mock-T controls – a common tumor adaptation mechanism [22]. These included the glycolysis-glucogenesis, pentose-phosphate, glycerophospholipid metabolism, glycosylphosphatidylinositol GPI anchor biosynthesis, glycosaminoglycan and other glycan degradation pathways (**Fig. 7a**). Immunosuppression indicated by upregulation of *Pdcd1* expression was also observed at this timepoint along with downregulation of the immune response *Cxcl1* gene and the angiogenesis gene *Ptgs2* (**Fig. 7b**). More importantly, CAR-T-treated tumors downregulated EMT and EMT-associated pathways compared to mock-T controls (**Fig. 7a, b**). These included the E2F targets (including the *E2f8* gene), TGF-β, Wnt and hedgehog signaling pathway (**Fig. 7a, b**). Since EMT can promote cell invasion and tumor metastasis, the downregulation of this pathway at this timepoint could potentially explain the reduced lung metastasis observed in our *in vivo* model.

All these findings suggest that CAR-T-treated tumors exhibited less proliferation on day 2 post-treatment which was followed by increased immune activation on day 7, followed by downregulation of EMT and upregulation of glycolysis on day 12 post-treatment when compared to mock-T controls. These findings highlight the complexity of tumor response mechanisms to CAR-T cell treatment and demonstrate that TME remodeling and dramatic metabolic changes commence very shortly following cell therapy affecting tumor growth trajectory and metastatic potential.

### Single cell RNA sequencing analysis shows tumor- and fibroblast-specific downregulation of EMT signatures in response to CAR-T cell treatment

To study quantitative and qualitative changes in each cell population of the TME, we used 10x Genomics single cell RNA FLEX sequencing (scRNA FLEX-seq). Three tumors from each of the treatment groups (CAR-T, mock-T and saline-treated mice) were harvested on day 12 post-treatment and the CD45^+^ and CD45^-^ cells from each tumor were column-sorted using magnetic beads. The cells were fixed and sequenced. Biological replicates of CD45^+^ or CD45^-^ cells were pooled together post-sequencing for downstream analysis.

Through unsupervised clustering, 6 CD45^-^ clusters were identified. Clusters 5 and 6 were combined into one cluster (cluster 5) due to their small size and were excluded from analysis. In the CD45^-^ population, clusters 0 and 1 were identified as major tumor clusters exhibiting high expression levels of epithelial cell markers (*Krt8, Krt18, Krt19*) (**Fig. 8a-c**). The proportion of tumor cluster 0 plus cluster 1 was observed lower in the A101 CAR-T-treated group, as compared to saline and mock-T-treated groups, indicating superior tumor clearance by CAR-T cells. At this timepoint no significant difference was observed in *Msln* expression between the treatment groups for cluster 0. However, there was a significant decrease of *Msln* in cluster 1 of the CAR-T-treated group compared to mock-T (pval < 0.05, log2FC = −0.529, pval adj. = 1, data not shown). Additionally, cluster 1 of the mock-T group significantly upregulated *Msln* compared to saline (pval < 0.05, log2FC = 0.49, pval adj. <0.05, data not shown). All these findings suggest target-specific effector function of the A101 CAR-T cells.

**Fig. 8.**
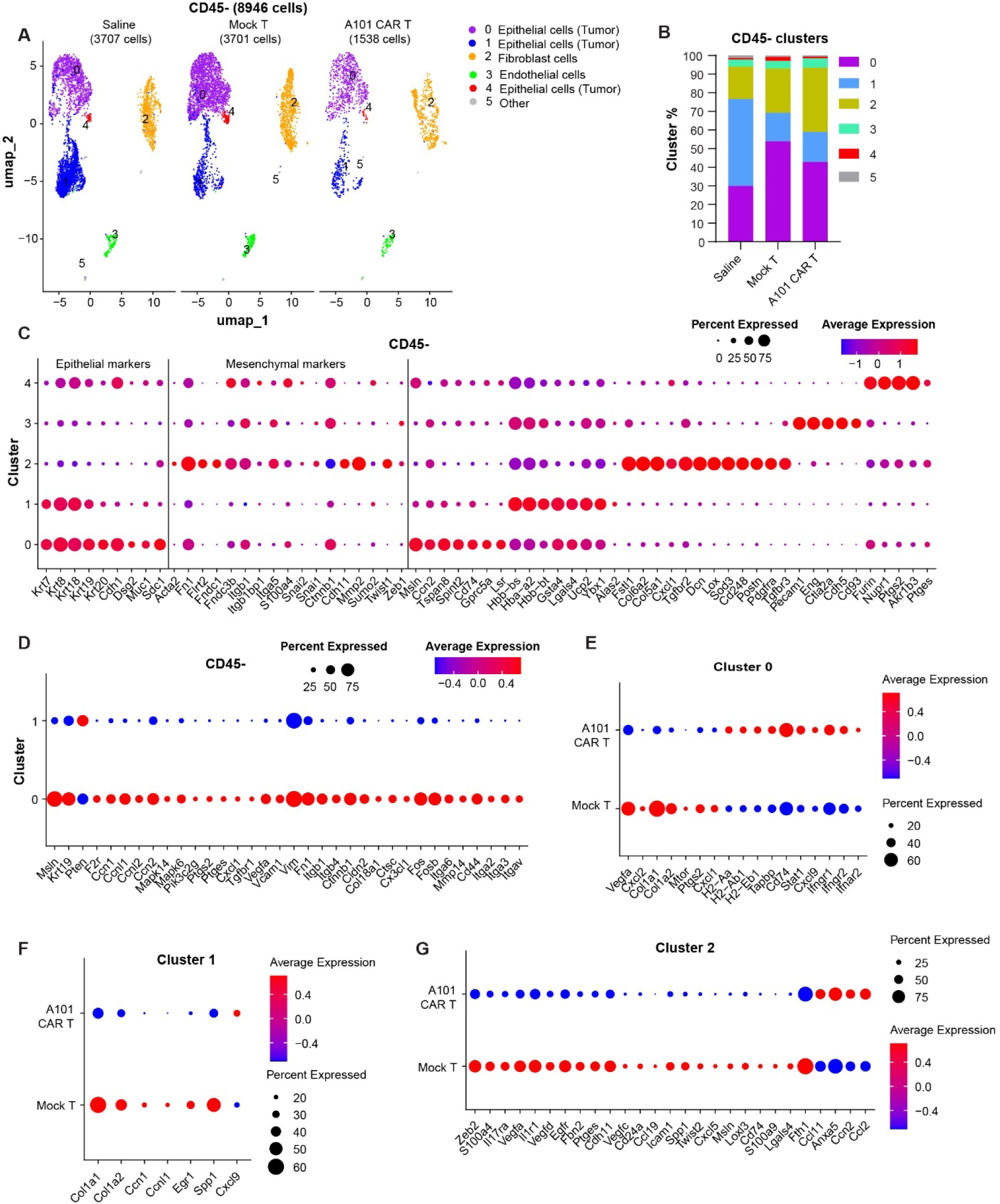
scRNA FLEX-seq identifies CD45^−^ cell populations in 344SQ tumors. 344SQ tumor-bearing 129sv mice were treated with one dose of 3 x 10^6^ A101 CAR-T or mock-T cells or saline. Tumors were harvested on day 12 post-administration of T cells for single cell RNA sequencing. n = 3. **a** UMAPs of CD45^-^ cell types sorted from 344SQ tumors. The clusters were color-coded based on their annotated cell names. **b** Relative quantification of cluster size in each treatment group. **c-g** Dot plots visualize gene expression in indicated clusters. Red indicates a higher average expression of the gene, and a larger dot size indicates that the gene is present in a larger percentage of cells within that cluster. **c** Scaled expression of cluster annotation genes. **d-g** Scaled expression of significant, differentially expressed genes in the indicated CD45^−^ clusters (p<0.05 for **e** and **f** and padj.<0.25 for **d** and **g**).

Comparison of cluster 0 to cluster 1 showed that cluster 0 had higher *Msln*, *Krt19*, *Ptgs2* and *Cxcl1* expression (**Fig. 8d**). In addition, cluster 0 tumor cells exhibited an increase in EMT genes when compared to cluster 1. These included: *Msln, Tgfbr1, Vegfa, Vcam1, Vim, Fn1, Itgb1, Itgb4, Ctnnb1, Cldn2, Col18a1, Ctsc, Cx3cl1, Fos, Fosb, Itga6, Mmp14, Cd44, Itga2, Itga3 and Itgav* (**Fig. 8d**).

Genes *Ptgs2, Cxcl1*, *Vegfa, Cxcl2, Col1a1* and *Col1a2* were downregulated in cluster 0 of the A101 CAR-T-treated tumor compared to the mock-T-treated group which are involved in angiogenesis and tumor-promoting inflammation as well as EMT and tumor progression (**Fig. 8e**) [23, 24] [25]. Additionally, upregulation of antigen presentation genes including *H2-Aa, H2-Ab1, H2-Eb1, Tapbp* and *Cd74* as well as *Stat1* and *Cxcl9* - a chemokine that recruits immune cells to the tumor, was observed in the CAR-T group (**Fig. 8e**) [26]. Considering that *Ifng* and *Ifna* can stimulate antigen presentation, this indicates that CAR-T cell treatment likely had a stronger effect in cluster 0 compared to mock-T cell treatment [27]. Increased expression of *Ifngr1, Ifngr2* and *Ifnar2* in cluster 0 of the CAR-T-treated group (compared to mock-T-treated), indicates this could be a tumor adaptive mechanism to Ifn-γ and Ifn-α signaling mediated by CAR-T cell therapy (**Fig. 8e**).

The other major tumor cell population identified, cluster 1, consists of *Msln^-/low^ Krt19^+^ Pten^+^* cells. Cluster 1 was decreased upon T cell treatment in both the A101 CAR-T and mock-T groups. However, the A101 CAR-T group showed upregulation of *Cxcl9* and downregulation of EMT-related genes such as *Col1a1* and *Col1a2* when compared to mock-T (**Fig. 8f**). *Cxcl9* is considered a positive prognostic marker along with *Pten* which is a tumor suppressor [26, 28]. Consistent with *Pten* upregulation, is the concomitant downregulation of *Pten* downstream targets such as *F2r*, *Ccn1*, *Ccnl1*, *Ccnl2*, *Ccn2*, *Mapk14*, *Mapk6* and *Pik3c2g* (**Fig. 8d**) [29]. This indicates that cluster 1 is potentially less aggressive tumor than cluster 0 and more sensitive to T cell mediated killing.

A proportional increase in cluster 2 fibroblasts was observed in the A101 CAR-T group compared to all other treatment groups. Fibroblasts have wound healing function, and their presence is indicative of stress [30]. Cluster 2 cells in the CAR-T group show increased levels of *Ccl11, Anxa5* and *Ccn2* compared to mock-T (**Fig. 8g**). These markers are associated with wound healing, tissue repair and response to injury and stress [31–33]. This indicates that damage has occurred in response to CAR-T cell treatment. In addition, there is an increase in *Ccl2* expression in the CAR-T group fibroblasts (**Fig. 8g**). *Ccl2* is responsible for attracting monocytes and other immune cells and could contribute to myeloid-mediated immunosuppression [34]. Fibroblasts from the CAR-T group also showed decreased expression of EMT-related markers including *Zeb2, Vegfa, Vegfd, Vegfc, Twist2, Cxcl5, Msln* and *Lgals4* (**Fig. 8g**).

In summary, both total RNA-seq and scRNA-seq data show downregulation of the EMT pathway, Cxcl1 and Ptgs2 in the A101 CAR-T-treated group compared to mock-T 12 days after treatment. At the same time, the reduction of the *Msln*^+^ tumor cluster 0 and the increased expression level of *Ccl2*^+^ in fibroblasts of the CAR-T-treated tumors could indicate a CAR-T specific response. The reduction of the less aggressive *Pten^+^* tumor cluster 1 relative to other tumor clusters could indicate a tumor selection mechanism in response to T cell treatment.

### A101 CAR-T-treated tumors upregulate *P2rx1^+^* neutrophils and downregulate *Cd274^+^ Lcn2^+^* neutrophils

Analysis of the CD45^+^ population using scRNA FLEX-seq, revealed changes in the myeloid and lymphoid cell compartment in the A101 CAR-T group (**Fig. 9a-d**).

**Fig. 9.**
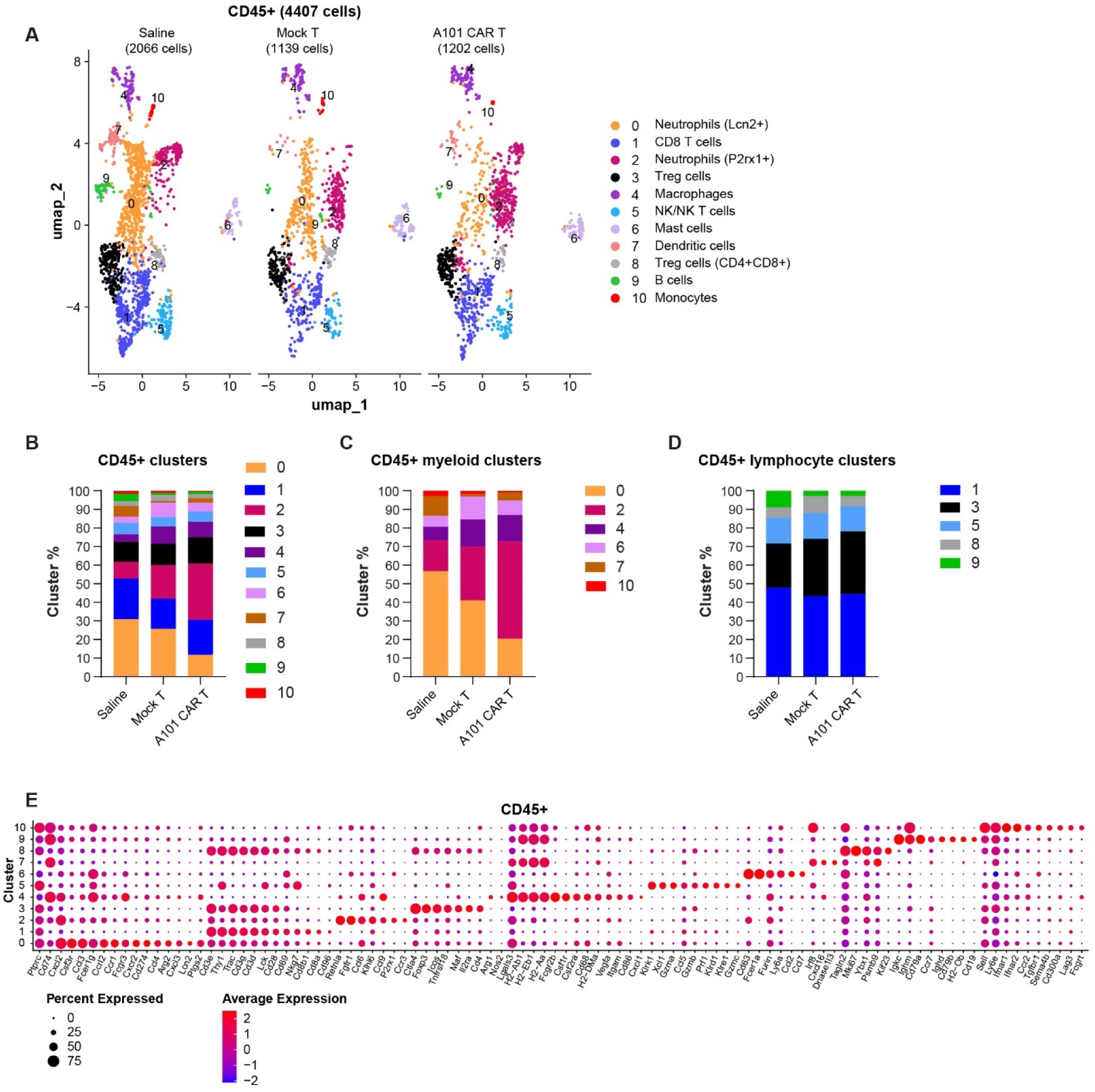
scRNA FLEX-seq identifies CD45^+^ cell populations in 344SQ tumors. **a** UMAPs of CD45^+^ cell types sorted from 344SQ tumors. The clusters were color-coded based on their annotated cell names. **b-d** Relative quantification of cluster size in each treatment group. **e** Dot plot visualizes gene expression in indicated clusters. Red indicates a higher average expression of the gene, and a larger dot size indicates that the gene is present in a larger percentage of cells within that cluster. Scaled expression of cluster annotation genes.

In the myeloid cell compartment, changes were observed in clusters 0 and 2 which were both identified as neutrophils. Specifically, cluster 0 was greatly reduced in the A101 CAR-T-treated group. These are *Ccl4^+^ Cd274^+^ Lcn2*^+^ *Ptgs2*^+^ immunosuppressive neutrophils implicated in EMT induction in tumor cells (**Fig. 9e**, **Fig. 10a**) [35–37]. This is consistent with the observed downregulation of the EMT pathway in the CAR-T group. Compared to mock-T, *Lcn2*^+^ neutrophils in the CAR-T group showed increased expression of survival (*Hspa1a, Socs1, E2f4, Eef2, Pdcd4*) and phagocytosis genes (*Maf, Nudt4, Clu, Gbp2, Isg15, Vamp2, Map3k8*) (**Fig. 10b**). They also downregulated *Lcn2* and markers associated with chemotaxis (*Ccrl2, S100a8, S100a11, S100a9, S100a4, Fpr2, Calm1, Ctla2a*) and tissue remodeling (*Ctsb, Ctsd*) (**Fig. 10b**). Collectively, this indicates that *Lcn2*^+^ neutrophils in the CAR-T group have a more phagocytic role and are less inflammatory.

**Fig. 10.**
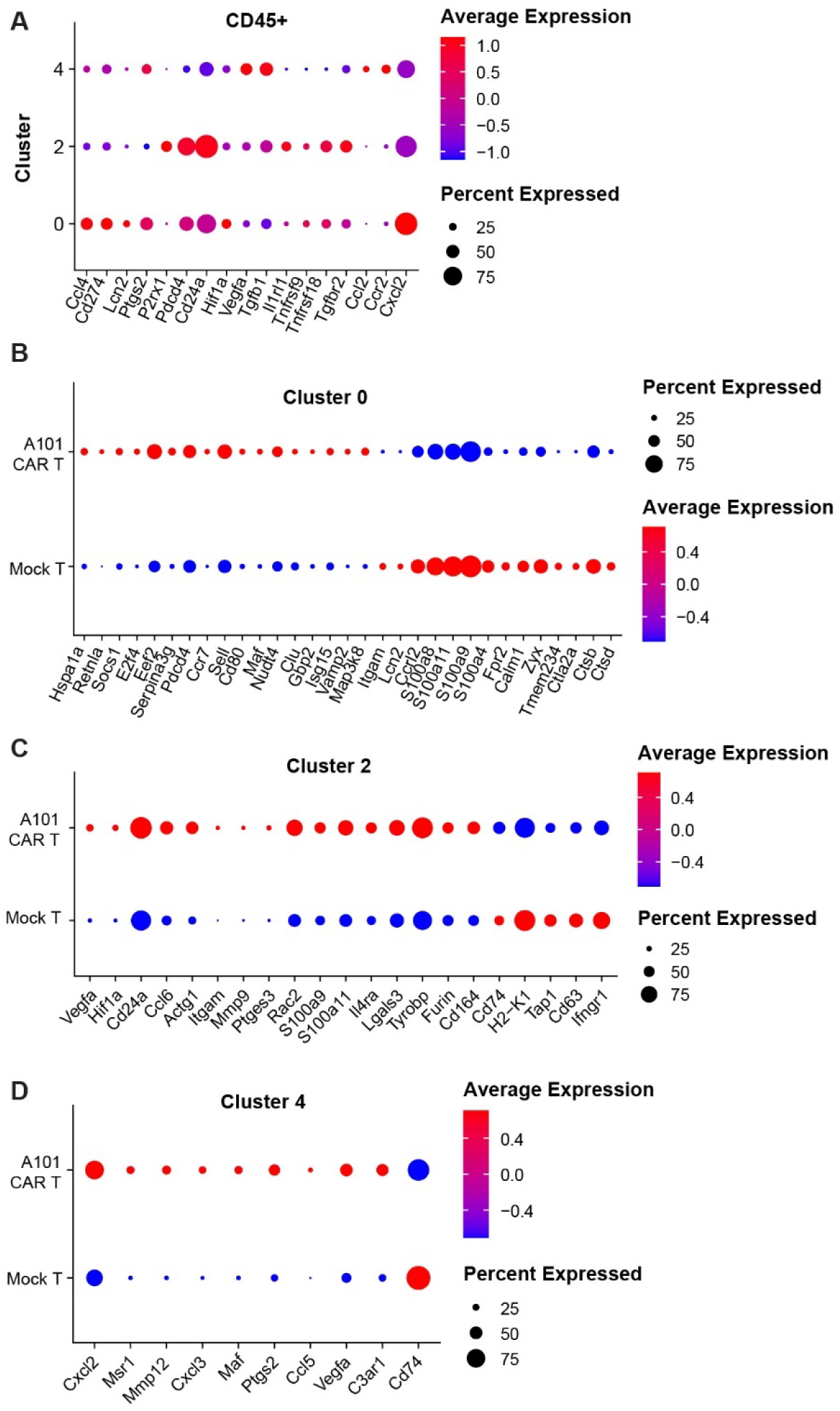
scRNA FLEX-seq identifies immunosuppressive myeloid cell populations in 344SQ tumors. **a-d** Dot plots visualize gene expression in indicated clusters. Red indicates a higher average expression of the gene, and a larger dot size indicates that the gene is present in a larger percentage of cells within that cluster. Scaled expression of significant, differentially expressed genes in the indicated CD45^+^ clusters (p<0.05 for **b**, **c** and **d** and padj.<0.25 for **a**).

In contrast, cluster 2 neutrophils were increased in the A101 CAR-T group compared to mock-T. These are *P2rx1^+^* neutrophils [38–40]. *P2rx1*^+^ neutrophils are associated with positive anti-tumor responses and inflammation [39]. Compared to mock-T, *P2rx1*^+^ neutrophils from the CAR-T group showed increased expression of *Vegfa, Hif1a, Cd24a, Ccl6, Itgam, Mmp9, Il4ra, Lgals3, Tyrobp,* and *Furin* (**Fig. 10c**).

An increase in cluster 4 was also observed in both the CAR-T and mock-T groups. These are *Ccl2^+^ Ccr2^+^ Cxcl2*^+^ macrophages that have been reported to inhibit immune cell responses by recruiting and inhibiting the differentiation of monocytes [34]. Cluster 4 macrophages in the CAR-T group showed increased expression of *Cxcl2, Cxcl3, Ptgs2, Ccl5* and *Vegfa* while they significantly downregulated *Cd74* compared to the mock-T group (**Fig. 10d**). This indicates a more tolerogenic phenotype in the CAR-T group compared to the mock-T control.

In the lymphoid cell compartment, two clusters of Treg cells were detected in the TME. Cluster 3 exhibited a classic Treg cell phenotype while cluster 8 was *CD4^+^ CD8^+^* and exhibited a more activated phenotype (**Supplemental Fig. 4b**). A slight increase in the Treg cell cluster 3 and a decrease in cluster 8 was observed in the CAR-T group compared to controls indicating a potential tumor adaptation mechanism.

Interestingly, the relative percentage of B cells (cluster 9) did not change between the CAR-T group and the mock-T control. However, the B cells in the CAR-T group showed a more activated phenotype (*Ighg1, Ccl3, Ccl6*) (**Supplementary Fig. 4c**).

Collectively, this data indicates the differential regulation of neutrophils in response to CAR-T cell therapy and potential immunosuppression mediated by *Cxcl2*^+^ macrophages in the TME. It also indicates an increase in Treg cells and activated B cells in a manner consistent with our flow cytometry and CIBERSORT analysis.

## Discussion

In this study we have engineered a nanobody-based CAR-T cell that is specific for mouse mesothelin. To generate A101 CAR-T cells, we chose to incorporate the camel-derived A101 sequence, and the human truncated EGFR sequence into a second-generation CAR construct. Mouse T cells exhibited good transduction efficiency rates with A101 CAR-T cells showing antigen specific effector function against a variety of mouse-derived tumor cell lines expressing Msln *in vitro*. Despite the high effector to target ratio required for the *in vitro* assays, mouse A101 CAR-T cells were able to reduce tumor growth rate in two separate syngeneic, subcutaneous models, after a single dose and in hosts that did not receive lymphodepletion. In the 344SQ model, a single dose of A101 CAR-T cells led to decreased lung metastasis and improved survival rates. This provided us with a powerful tool to investigate the effects of CAR-T cell therapy in the TME of metastatic solid tumors over time without confounding factors such as lymphodepletion and multiple dosing.

Tumor profiling using flow cytometry, showed that A101 CAR-T cells homed to the tumor within 48 hours after inoculation and induced changes in the TME that could be observed 12 days post-treatment in unprimed hosts. Most notably, Treg and B cells increased over time in all treatment groups with a higher increase observed in the A101 CAR-T-treated group compared to mock-T control. This was consistent with CIBERSORT data which also showed an increase in Treg cells and plasma cells in the same treatment group. scRNA-seq analysis also showed a proportional increase in Treg cells and a more activated B cell phenotype in the CAR-T-treated group compared to the mock-T control. While Treg cells are typically associated with immunosuppression and tumor-promoting effects, the role of B cells in the TME is less clear. Total RNA-seq and scRNA-seq analysis showed increased B cell activation on day 12 post-treatment in the CAR-T-treated group relative to mock-T control. B cells can have a positive anti-tumor effect through antigen presentation, the release of tumor-reactive antibodies, antibody-dependent cellular cytotoxicity and complement-dependent cytotoxicity [42]. B cell differentiation into tumor reactive plasma cells has been positively associated with improved melanoma and lung cancer patient survival [42]. However, B cells can also have immunosuppressive functions through the secretion of IL-10 and TGF-β [43]. In our studies, no significant differences were observed in *Il-10* and *TGFb* expression by B cells when the CAR-T and mock-T treatment groups were compared (data not shown). The only difference observed was that B cells in the CAR-T-treated group exhibited a more activated phenotype including upregulation of *Ighg1* expression. The significance of B cell activation and upregulation of *Ighg1* in the TME of 344SQ tumors remains unclear and it needs further investigation.

Another gene that was differentially expressed in the CAR-T group was *Ptgs2* (also known as *Cox-2*). Downregulation of this gene was observed using total RNA-seq two days and 12 days post-treatment. scRNA-seq also revealed the downregulation of *Ptgs2* in the predominant tumor cluster 0 and an overall decrease in the immunosuppressive *Cd274*+ *Lcn2+ Ptgs2*+ neutrophil cluster 0 in the CAR-T-treated group. *Ptgs2* is associated with tumor immune evasion by regulating myeloid derived suppressor cells [44, 45]. Studies have shown that Ptgs2 expression in gastric cancer patients is associated with poor prognosis [44]. The downregulation of this gene in the CAR-T-treated tumor transcriptome we have observed using both total and scRNA-seq is in accordance with our observation of improved survival rates in mice treated with A101 CAR-T cells.

Signaling pathway analysis of tumor RNA at different timepoints revealed rapid metabolic changes in the TME. Specifically, EMT was upregulated on day 7 post-treatment and downregulated on day12 in the CAR-T-treated group compared to mock-T. Interestingly, day 7 was the timepoint where EMT upregulation coincided with upregulation of Kras signaling which is known to activate EMT in tumors [46]. Inflammation, transendothelial leukocyte migration and immune activation pathways were also upregulated at this timepoint along with ECM remodeling - presumably to facilitate immune cell recruitment. Studies have shown that EMT and inflammation can drive fibroblast generation and tissue fibrosis [47]. This is consistent with the proportional increase in fibroblasts and the increased levels of *Ccl2* expression in these fibroblasts observed on day 12 post-treatment in the CAR-T-treated group using scRNA-seq. This indicates that fibroblast generation started around day 7 post-treatment. Fibroblasts have been associated with immunosuppression through the recruitment of pathogenic myeloid cells [31]. It has been previously shown that targeting fibroblasts along with mesothelin in fibrotic pancreatic tumor models improves CAR-T cell persistence and host survival [11]. Since we observed a proportional increase in fibroblasts in the TME, it is possible that targeting fibroblasts along with Msln may enhance tumor response in our 344SQ model too. However, studies have also shown that certain fibroblast types such as *Ccl19^+^* fibroblasts, can have anti-tumor function potentially by supporting CD4^+^ T and B/plasma cell function [48]. In our data, *Ccl19* was significantly downregulated in fibroblasts of the CAR-T-treated group compared to mock-T control. The role of fibroblasts in the TME can be multifaceted and further investigation of the identified population is required. Additionally, studies have shown that a tumor adaptation mechanism exists whereby the Ccl2/Ccr2/Cxcl2 axis promotes the recruitment of monocytes which in turn can acquire pro-tumorigenic phenotypes and contribute to immunosuppression [34, 49]. This is consistent with our finding of decreased antigen presentation (*Cd74*) and increased *Cxcl2* expression in macrophage cluster 4 of the CAR-T-treated group.

In summary, our data indicate that CAR-T cell treatment led to increased tumor damage which in turn stimulated inflammation and a type II EMT response which is associated with fibroblast generation and wound healing [47]. Contrary to type II EMT response, type III EMT response is associated with tumor cell invasiveness and metastasis [47, 50]. According to both total RNA-seq and scRNA-seq data, EMT was significantly downregulated on day 12 post-treatment in the CAR-T-treated tumors compared to mock-T which is consistent with our observation of improved survival and reduced lung metastases in CAR-T-treated mice. This lends support to our hypothesis that the EMT response observed on day 7 post-treatment is a type II EMT response further highlighting the importance of timing in understanding TME dynamics.

In accordance with the reduction in EMT observed on day 12 post-treatment, *Lcn2*^+^ neutrophils which have been reported to promote EMT, were also downregulated in CAR-T-treated tumors [35–37]. *Lcn2*^+^ neutrophils were found to express *Ccl4* and *Cd274* and this phenotype has been previously described to exert immunosuppression via tumor associated macrophage (TAM) recruitment through *Ccl4* and T cell inhibition via *Cd274* signaling [36]. In addition to downregulation of the immunosuppressive EMT-promoting *Lcn2*^+^ neutrophils, CAR-T cell-treated tumors showed upregulation of *P2rx1*^+^ neutrophils. *P2rx1*^-^ *Nrf2*^+^ neutrophils have been shown to have immunosuppressive, pro-tumorigenic function in clinical and animal studies of metastatic pancreatic cancer [39]. Therefore, it is possible that the increase in *P2rx1*^+^ *Nrf2*^-^ neutrophils in our model could have a beneficial effect for the host which is consistent with the decreased primary tumor growth rate and reduced metastasis observed *in vivo*. Depletion of neutrophils and/or TAMs in combination with A101 CAR-T treatment could help determine whether the observed differential regulation of these myeloid populations is due to a direct or indirect effect of the CAR-T cell treatment and demonstrate their importance in tumor progression in future studies.

In summary, using flow cytometry, total RNA-seq and scRNA FLEX-seq to profile the TME in immunocompetent hosts, our study highlights the complexity of tumor responses to cell therapy and indicates that even when CAR-T cells do not persist, they can induce significant changes in the TME that alter tumor growth trajectory and metastatic potential. This study also highlights potential targets such as *Lcn2^+^* neutrophils and *Ccl2*^+^ fibroblasts for therapeutic intervention to enhance anti-mesothelin CAR-T cell therapy for metastatic solid tumors.

## Methods

### Animals

All animal experiments were approved by the NCI-Bethesda Animal Care and Use Committee (ACUC protocol no.: TGOB-001) and were carried out in accordance with the guidelines. *129S2/SvPasCrl, C57BL/6NCrL* and *B6.SJL-PtprcaPepcb/BoyCrCrl* mice were purchased from Charles River Laboratories. Mice were housed in a specific pathogen-free (SPF) condition at ambient temperature (22 ± 2 °C), air humidity 40–70% and 12 h dark/12 h light cycle and had free access to water and chow. Animal health status was routinely checked by qualified veterinarians.

For the transplant models, tumor cells were inoculated into female mice aged 8-10 weeks of similar weight, randomized before tumor inoculation. All animal experiments were performed in the same well-controlled pathogen free facility. No tumors exceeded the maximum diameter of 2 cm approved by ACUC at the time the mice were sacrificed for tumor analysis or in the survival studies.

### Cell lines

AB12, Panc02, A431, A431/H9, KLM1, KLM1 MSLN KO, 344SQ, AE17, 531LN2 and LLC cells were used in this study (sources listed in **Supplementary Table 1**). The cell lines 344SQ and 531LN2 were kindly provided by Dr. Jonathan M Kurie as a gift from MD Anderson Cancer Center. TC-1 cells were kindly gifted by Professor TC Wu (JHU). GFP^+^ PDAC 95575 and GFP^+^ CREP133239 were kindly given by Dr. Mitchell Ho (NIH). GFP^-^ PDAC 95575 cells were kindly gifted by Dr. Serguei Kozlov (NIH). All cell lines were tested for the presence of mycoplasma contamination (MycoAlert Mycoplasma Detection Kit, Lonza). All cancer cells were cultured at 37 °C with 5% CO2 in media supplemented with 10% Fetal Bovine Serum (FBS), 100 U/ml penicillin-streptomycin and L-glutamine. 531LN2, 344SQ, AE17, AB12, TC-1, Panc02, CREP133239 and all human cell lines were cultured in RPMI media. LLC and PDAC 95575 cells were cultured in DMEM media. Mouse T cells were cultured at 37 °C with 5% CO2 in DMEM (RPMI was used for human T cells) including 10% FBS, 1mM sodium pyruvate, 0.1mM non-essential amino acids, 50U/ml penicillin-streptomycin, L-glutamine and 50μM 2-b-mercaptoethanol.

### A101 epitope binding assay

Maxisorp 96-well plate was coated with 5 µg/ml mesothelin fragments overnight at 4°C. The plate was washed with PBST three times and pre-blocked with 100% superblock for two hours. The plate was washed with PBST three times and 5 μg/ml A101 V_H_H in PBST and 10% superblock was added to the plate for one hour. The plate was then washed six times with PBST. 1/5000 anti-FLAG HRP in PBST and 10% superblock was added for one hour. Plate was washed 6 times and developed with TMB substrate. Finally, 0.25M sulphuric acid was added and the plate was read using a spectrophotometer (Molecular Devices) at 450nm.

### CAR-T cell generation

The V_H_H domain of A101 was fused with a mouse CD8a hinge domain followed by a mouse CD8a transmembrane (TM) domain (Uniprot reference sequence mouse P01731) and the mouse 4-1BB (Uniprot reference sequence mouse P20334) and CD3z (Uniprot reference sequence mouse P24161) signaling domains. The CAR sequence in the plasmid was expressed under the EF1a promoter and the sequence was codon optimized for expression in the mouse T cell. This CAR targets both mouse and human mesothelin. The plasmid was packaged into a lentivirus and used to generate the mouse CAR-T cells. Viral titers were determined in vitro using HEK293 T cells prior to T cell transduction.

Mouse T cells were isolated from the spleen using T cell isolation kit (Miltenyi) according to the manufacturer’s instructions. T cells were stimulated with αCD3/αCD28 Dynabeads (Gibco) at 1:1 bead to cell ratio in the presence of rhIL2 (50 IU/ml) for 24 hrs. Cells were then transduced at MOI of 1.2 for 24 hrs in the presence of lentiblast (1:1000 dilution, OZ Biosciences).

For human CAR-T cell generation, PBMCs from healthy donors were provided by the NIH Clinical Center Department of Transfusion Medicine as part of their IRB approved and consented healthy donor program. PBMCs were stimulated with αCD3/αCD28 Dynabeads (Gibco) at 2:1 bead to cell ratio in the presence of rhIL2 (100 IU/ml) for 72 hrs. Cells were then transduced using a lentiviral vector containing the human A101 or YP218 CAR construct at MOI of 5. Cells were spinoculated at 1000xg for 2 hrs at 32°C and cultured in media containing 10 μg/ml protamine sulfate for 24 hrs (Thermo Fisher Scientific). Transduction efficiency was calculated based on expression of CAR assessed by staining with APC anti-V_H_H (GeneScript) or PE anti-EGFR (R&D Systems) expression using flow cytometry.

### Cytotoxicity and cytokine release assay

For checking the cytotoxicity of A101 CAR-T cells *in vitro*, *344SQ.GFP/Luc, AE17.GFP/Luc, 531LN2.GFP/Luc*, *TC-1.GFP/Luc, A431.GFP/Luc*, *A431/H9.GFP/Luc*, *KLM1.GFP/Luc* and *KLM1 MSLN KO.GFP/Luc* cells were co-cultured with un-transduced (mock-T) or A101 CAR-T cells at the E:T ratios of 0, 0.2, 0.4, 0.8, 1.6, 3, 6, 12.5, 25 and 50, in triplicates, in 96-well flat-bottomed plates. Luciferase activity from cell lysates was determined using the Luciferase Assay System (Promega) by measuring relative light unit (RLU). Target cells alone (E:T = 0) were set as negative control to detect spontaneous death. The percentage of killing was calculated using the formula: % killing = (1-(tested sample RLU/spontaneous death RLU))*100. After 24 hours of coculture, supernatants containing the effector T cells from the E:T = 50 ratios were collected and analyzed for IL-2, IFN-γ and TNF-α expression using intracellular flow cytometry. IFN-γ levels were also assessed in supernatants using the Quantikine ELISA Kit (R&D Systems) according to the manufacturer’s instructions. For checking the cytotoxicity of A101 CAR-T cells against the *PDAC 95575.GFP/Luc* and *CREP133239.GFP/Luc* cells, target cells were co-cultured with mock-T or A101 CAR-T cells at E:T ratios 0, 3, 6 and 12, in triplicates, in 96-well flat-bottomed plates. GFP levels indicating target cell survival were measured at 2-hour intervals for a total of 78 hours using IncuCyte live cell imaging (Sartorius). Target cell survival is expressed as the total surface area occupied by GFP^+^ target cells normalized to surface area occupied by GFP^+^ target cells at 0 hour.

### In vivo models

For the syngeneic subcutaneous tumor models, 1 × 10^6^ 344SQ and PDAC 95575 cells were suspended into 100 ul PBS and inoculated s.c. into the right flank of *129S2/SvPasCrl* and *C57BL/6NCrL* mice respectively. Tumor size was measured two to three times per week using caliper. Tumor volume was calculated as width×width×length×0.4 and randomly assigned to the various treatment groups. In all experiments, the total number of T cells injected was equal between the CAR-T and mock-T-treated groups.

For the 344SQ model, mock-T or A101 CAR-T cells were i.v. injected into tumor-bearing *129S2/SvPasCrl* mice at the dose indicated in the figure legend when tumor volume reached 100–150 mm^3^; saline was used as control. For flow cytometry exepriments, tumors were collected on day 2, 7 and 12 post-T cell administration. Blood was collected at day 2 and 7 post-T cell administration. For the total RNA-seq experiment, tumors from the CAR-T-treated or mock-T-treated mice were harvested on day 2, 7 and 12. For the single cell RNA FLEX sequencing experiment, tumors from the CAR-T-treated, mock-T and saline-treated mice were harvested on day 12. For all survival studies, mice were euthanized when tumor volume reached 1,200 mm^3^.

For the PDAC 95575 model, mock-T or A101 CAR-T cells from *B6.SJL-PtprcaPepcb/BoyCrCrl* mice were i.v. injected into tumor-bearing *C57BL/6NCrL* mice at the dose indicated in the figure legend when tumor volume reached 100–150 mm^3^; saline was used as control. Tumors were harvested on day 2 and 15 post-treatment and analyzed by flow cytometry.

### Flow cytometry

Tumor tissue was dissected and digested with 0.25 mg/mL Liberase (Sigma) plus 0.5 mg/mL DNase I (Sigma) in DMEM media (Thermo Fisher) for 30 min at 37 °C. Digestion mixture was passed through 100 μm cell strainer (FALCON) to prepare single cell suspension and washed with stain buffer (PBS, GIBCO with 5% FBS, Lonza). Blood was collected through submandibular bleeding and red blood cells (RBC) were removed using the RBC lysis buffer (BioLegend). After washing with PBS, cells were collected. In certain experiments, live cells from tumor tissue were column sorted prior to flow cytometry according to manufacturer’s instructions (Miltenyi).

Unless otherwise indicated, DAPI (Invitrogen) or ZombieAqua (Invitrogen) was used for live/dead cells staining. Single cells were blocked with anti-mouse CD16/CD32 (Invitrogen) for 15 min followed by cell surface staining with antibodies diluted in stain buffer (BD Biosciences) for 30 min on ice. For detecting intracellular cytokine, single cell suspensions were stained for IL-2, IFN-γ and TNF-α using the Fixation/Permeabilization Solution Kit (BD Biosciences). Foxp3 staining was performed using the Foxp3/Transcription Factor staining buffer set (eBiosciences) according to the manufacturer’s instructions. The list of antibodies used for flow cytometry can be found in **Supplementary Table 1**. Flow cytometry was performed on a CytoFLEX flow cytometer (Beckman Coulter, USA) and analyzed using FlowJo software (FlowJo LLC, USA).

### Immunohistochemistry staining

Tumors were fixed in 10% formalin (Sigma) overnight at 4°C. Paraffin-embedding, H&E staining and immunohistochemical staining for Msln and CD3 was performed by the Molecular Histopathology Laboratory (NCI at Frederick, MD). The Bond Polymer Refine Kit (Leica Biosystems, Cat# DS9800) was used along with the following antibodies: anti-mouse Msln (1:1000, Invitrogen, Cat# PA5-79698), CD3 (1:100, Bio-Rad Laboratories, Clone CD3-12, Cat# MCA1477) and Rabbit Anti-Rat (1:100, Vector Laboratories, Cat# BA-4001). Images were obtained using an Aperio dgital slide scanner (Leica Biosystems) and analysed using Aperio ImageScope (Leica Biosystems) and Halo software (Indica Labs).

### Total RNA sequencing

Tumor tissue was stored in RNAlater Stabilization Solution (Invitrogen), homogenized and RNA was isolated using RNAeasy kit (Qiagen) according to the manufacturer’s instructions. Samples were pooled and sequenced on NovaSeq Xplus 1.5B using Illumina® Stranded Total RNA Prep, Ligation with Ribo-Zero Plus and paired-end sequencing. The samples had 87 to 174 million pass filter reads with more than 95% of bases above the quality score of Q30. Reads of the samples were trimmed for adapters and low-quality bases using Cutadapt before alignment with the reference genome (mm10) and the annotated transcripts using STAR. The average mapping rate of all samples was 91%. Unique alignment was above 72%. There were 6.46 to 15.04% unmapped reads. The mapping statistics were calculated using Picard software. The samples had 1.75% ribosomal bases. Percent coding bases were between 30-42%. Percent UTR bases were 28-40%, and mRNA bases were between 61-80% for all the samples. Library complexity was measured in terms of unique fragments in the mapped reads using Picard’s MarkDuplicate utility. The samples had 54-73% non-duplicate reads. The gene expression quantification analysis (normalized count and raw count) was performed for all samples using STAR/RSEM tools. Downstream analysis and visualization were performed within the NIH Integrated Analysis Platform (NIDAP) using R programs developed by a team of NCI bioinformaticians on the Foundry platform (Palantir Technologies). RNA-Seq FASTQ files were aligned to the reference genome (mm10) using STAR [51] and raw counts data produced using RSEM [52]. The gene counts matrix was imported into the NIDAP platform, where genes were filtered for low counts (<1 cpm) and normalized by quantile normalization using the limma package [53]. Differentially expressed genes were calculated using limma-Voom [54]. Pre-ranked GSEA was performed using the fgsea package [55] and pathway enrichment analysis was performed using Fisher’s Exact Test [56]. Genes associated with pathways of interest from this analysis were visualized in heatmaps.

For the pathaway activity heatmap, Gene Set Variation Analysis (GSVA) was utilized to compute pathway enrichment scores from log-normalized and quantile normalized gene expression values of the samples [57]. These scores were then averaged based on the treatment groups to which the samples belonged. The enrichment scores for the CAR-T samples on days 2, 7, and 12 were calculated as ratios relative to the scores of the mock-T samples for comparison. These ratios were visualized using the R ComplexHeatmap package to illustrate the changes over time.

For the CIBERSORT analysis, we utilized the previously published LM22 signature matrix [21] of immune cells from human PBMCs. The matrix was deconvoluted and employed to impute cell fractions in our non-log normalized total RNA-seq data following the guidelines outlined on the CIBERSORTx website. To account for variations between the reference signature matrix and our dataset, we applied bulk (B-mode) batch correction. We opted to disable quantile normalization and selected 100 permutations to compute a null distribution for significance testing. The resulting cell fractions were calculated in relative values, allowing for comparisons across different samples [58].

### Single cell RNA FLEX sequencing

344SQ tumors from CAR-T-, mock-T- and saline-treated mice were harvested on day 12 post-treatment. Tumors from 3 different mice were harvested per treatment group. Tumor tissue was harvested in cold DMEM media supplemented with 10% FBS. Tissue was dissected and digested with 0.25 mg/mL Liberase (Sigma) plus 0.5 mg/mL DNase I (Sigma) in DMEM media (Thermo Fisher) for 30 min at 37 °C. Digestion mixture was passed through 100 μm cell strainer (FALCON) to prepare single cell suspension and washed with cold media. Dead cells were removed (Miltenyi kit) and CD45^-^ and CD45^+^ cells were column sorted using Miltenyi kits according to manufacturer’s instructions. Samples were fixed using the Chromium Next GEM Single Cell Fixed RNA Sample Preparation Kit (10x Genomics) in preparation for 10x Genomics singe cell RNA-FLEX sequencing according to the manufacturer’s instructions. Samples were stored at −80 °C until they were ready for sequencing.

Samples were thawed, pelleted, and resuspended in 0.5X PBS + 0.02% BSA for cell counting on the LunaFX7 and probe hybridization. Sample quality was determined based on single cellular suspension and homogeneity of cellular solution. A series of wash cycles and normalization pooling was performed to allow for 4×4 multiplexing. Capabilities of scFLEX 4-Plex include 8 capture lanes with 4-plex capture and an expected target recovery of 20,000 cells per lane (5,000 cells per sample). A total of 32 samples were 4×4 multiplexed and the pools were then loaded in 8 capture lanes according to the 10X Genomics Chromium Fixed RNA Profiling User Guide. 14 of these samples were not relevant to this study and therefore are not presented. The rest of the samples (18 samples) were subsequently analysed. These consisted of 3 biological replicates of CD45^-^ and 3 biological replicates of CD45^+^ cells per treatment group (CAR-T, mock-T, saline). Cell partitioning completed successfully with uniform emulsion consistency and the reverse transcription PCR was run overnight. The pre amplification and library construction were completed successfully. Single-cell suspension and droplet-based sequencing was performed. Three NovaSeq runs for FRP were performed. Base calling was performed using RTA 3.4.4, demultiplexing was performed using cellranger v7.2.0 (Bcl2fastq 2.20.0), and alignment was performed using cellranger v7.2.0 (STAR 2.7.2a). Sequenced reads were aligned to the 10x Genomics provided mouse reference sequence (refdata-gex-mm10-2020-A) and probe set (Chromium_Mouse_Transcriptome_Probe_Set_v1.0.1_mm10-2020-A).

Biological replicates of CD45^-^ or CD45^+^ samples were pooled together post-sequencing and analyzed. Seurat (version 4.4.0) was used to analyse single cell RNA-seq data [59]. The cells with UMI counts lower than the bottom 5% or higher than the top 5%, as well as those with more than 20% mitochondrial genes, were considered low quality and filtered out. We used scDblFinder to remove doublets. Markers of each main cell cluster were identified through the FindAllMarkers function. Differential analysis was performed using the FindMarkers function in Seurat (version 4.4.0) with test.use=’MAST’. Cluster-based filtering was performed to exclude *Ptprc*^+^ and *Ptprc*^-^ cells from the CD45^-^ and CD45^+^ samples respectively. Re-clustering was performed followed by manual annotation and differential gene expression analysis. We used clustering resolution of 0.2 for the CD45^-^ population and 0.3 for the CD45^+^ population. Pct threshold was set to 0.1.

### Statistics and reproducibility

N number of mice for each treatment group is indicated in each figure. For *in vitro* experiments, cells, or tissues from each animal were processed in biological triplicates minimum unless otherwise indicated. All data were shown as average ± S.E.M. Statistical analysis between two groups was conducted with a 2-tailed Student t test. Multiple comparisons were performed by using two-way ANOVA analysis with Tukey’s or Sidak’s multiple-comparison. Tumor growth curve analysis was conducted with two-way ANOVA (mixed-model) with Tukey’s multiple comparison. Kaplan-Meier curves were used to analyze the survival data. All statistical analysis was performed using GraphPad Prism 9 and the results of statistical analyses for each experiment are clearly indicated in the respective figures and in Figure Legends. p values < 0.05 were considered significant.

## Data Availability

Data are available within the Article and Supplementary Information.

## Acknowledgements

The cell lines 344SQ and 531LN2 were kindly provided by Dr. Jonathan M Kurie as a gift from MD Anderson Cancer Center. The TC-1 cell line was a kind gift from Professor. TC.Wu, Department of Pathology, SOM, JHU (NCI MTA. #49891-22). The GFP/Luc negative PDAC 95575 cell line used for the animal experiments was a kind gift from Dr. Serguei Kozlov, CAPR, NIH. This work was supported by the Intramural Research Program of the NIH, National Cancer Institute, Center for Cancer Research. We thank Dr. Brad Gouker and Dr. Baktiar Karim from the Molecular Histopathology Laboratory (MHL) for assistance with PFPE sections, immunohistochemistry staining and image analysis. We thank Dr. Geneti Ayana Gaga for his assistance with animal injections. The authors thank the Center for Cancer Research (CCR) Genomics Core, Bethesda, Maryland, the CCR Sequencing Facility, Frederick, Maryland and the CCR Collaborative Bioinformatics Resource, Bethesda, Maryland for assistance with single cell RNA FLEX and total RNA sequencing. This work used the computational resources of the National Insitutes of Health (NIH) HPC Biowulf cluster.

## Author contributions

Conceptualization: C.S., Q.J., R.H.; methodology: C.S., M.Z., Q.J., J.H., J.Z., R.H.; formal analysis: C.S.; investigation: C.S., Q.J., R.H.; resources: M.H., R.H.; data curation: C.S., M.Z., Q.J., J.B., R.H.; writing—original draft: C.S.; writing—review & editing: C.S., Q.J., M.H., M.Z., J.B., R.H.; visualization: C.S., M.Z., J.B.; study supervision: R.H.; funding acquisition: R.H..

## Additional information

Data are available within the Article and Supplementary Information.

**Correspondence** and requests for materials should be addressed to Raffit Hassan.

