## Supplementary Figures for "Temporal mapping of the anti-tumor effects of nanobody-based MSLN.CAR-T cell therapy in metastatic solid tumors"

Supp fig 1.

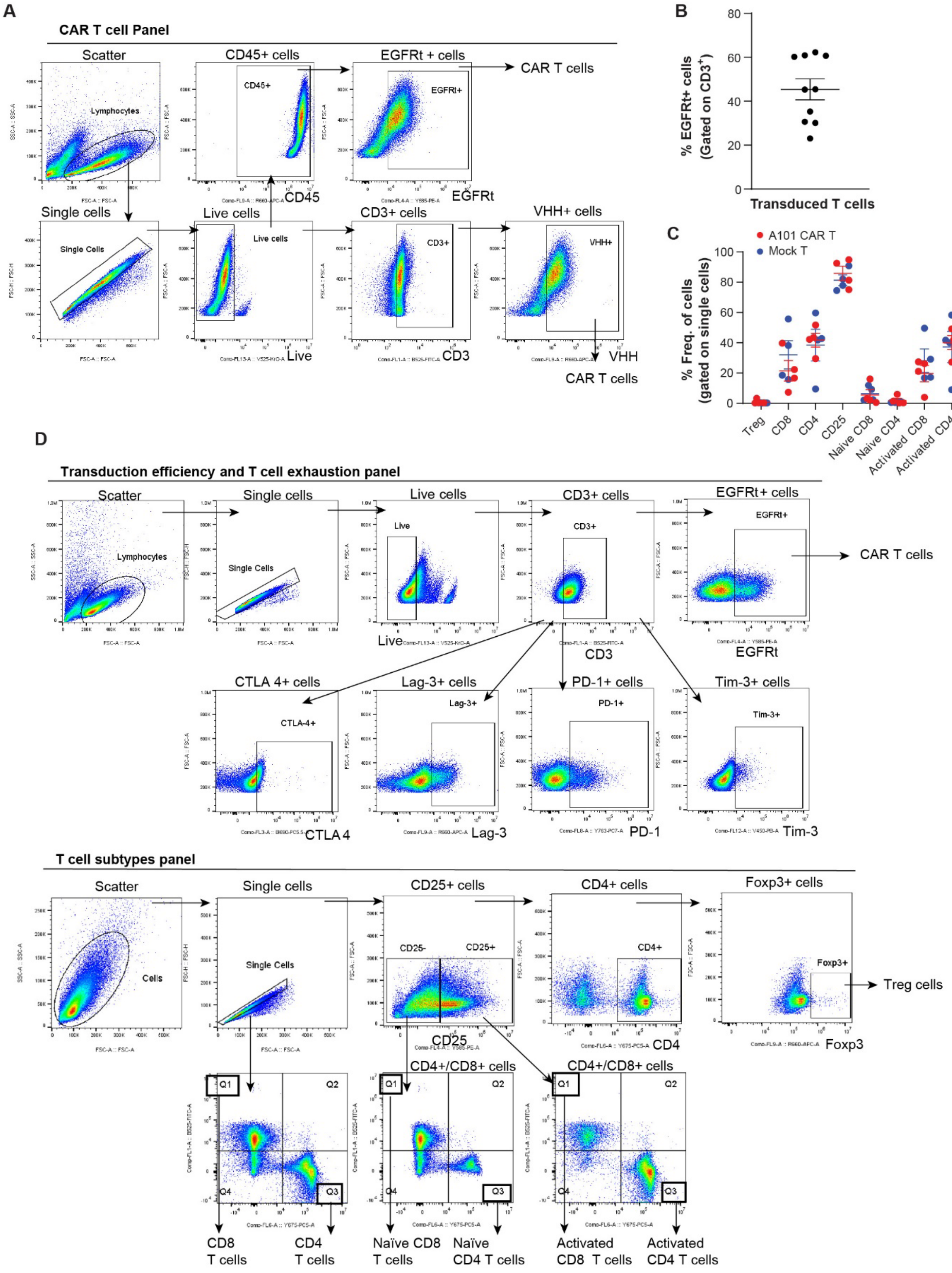
A
CAR T cell Panel
Scatter
Lymphocytes
CD45+ cells
CD45+
CD45
EGFRt + cells
EGFRt+
EGFRt
CAR T cells
Single cells
Single Cells
Live cells
Live
CD3+ cells
CD3+
CD3
VHH+ cells
VHH+
VHH
CAR T cells
B
% EGFRt+ cells (Gated on CD3+)
Transduced T cells
C
A101 CAR T
Mock T
% Freq. of cells (gated on single cells)
Treg
CD8
CD4
CD25
Naive CD8
Naive CD4
Activated CD8
Activated CD4
D
Transduction efficiency and T cell exhaustion panel
Scatter
Lymphocytes
Single cells
Single Cells
Live cells
Live
CD3+ cells
CD3+
CD3
EGFRt+ cells
EGFRt+
EGFRt
CAR T cells
CTLA 4+ cells
CTLA-4+
CTLA 4
Lag-3+ cells
Lag-3+
Lag-3
PD-1+ cells
PD-1+
PD-1
Tim-3+ cells
Tim-3+
Tim-3
T cell subtypes panel
Scatter
Cells
Single cells
Single Cells
CD25+ cells
CD25-
CD25+
CD25
CD4+ cells
CD4+
CD4
Foxp3 cells
Foxp3+
Foxp3
Treg cells
CD4+/CD8+ cells
CD4+/CD8+ cells
Q1
Q2
Q3
Q4
CD8 T cells
CD4 T cells
Naïve CD8 T cells
Naïve CD4 T cells
Activated CD8 T cells
Activated CD4 T cells

**Supplementary Fig. 1. A101 CAR-T cell product characterization *in vitro*. a** Gating strategy for determining transduction efficiency. **b** Transduction efficiency at MOI 1.2 ($n = 10$ independent samples). Data points are mean ± SE. Gating strategy is shown in **d**. **c** The percentages of indicated immune cell subpopulations in total single cells quantified based on multi-parametric flow cytometry analysis on day 6 post-transduction. Data points are mean ± SE ($n = 4$ per group) and groups were compared using two-way ANOVA with Sidak's multiple comparisons tests (from left to right; $F=0.09500$, 1.286, 0.4011, 0.5651, 0.1145, 0.1430, 1.039, 0.3552, $df=48$ for all). No significant differences were found ($p \geq 0.05$). Gating strategy is shown in **d**.

Supp fig 2.

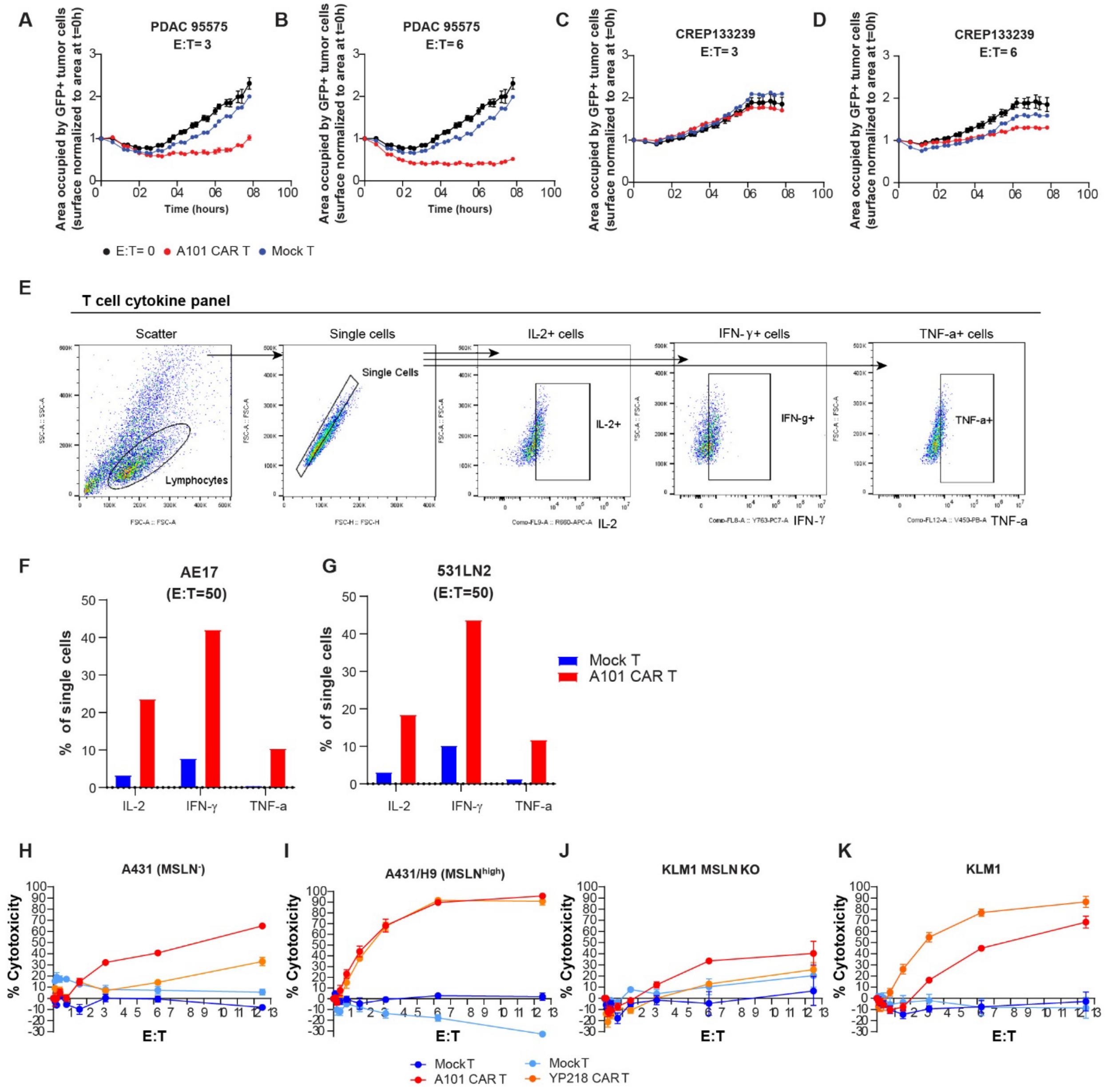


**Supplementary Fig. 2. Cytotoxic activity of A101 CAR-T cells *in vitro*. a-d** Cytotoxic activity of T cells during 78hrs of co-culture with GFP+ Msln+ murine tumor cell lines PDAC 95575 and CREP133239 at E:T ratio of 3 and 6 (n=3). Mean ± SE is shown. Groups were analyzed using two-way ANOVA with Tukey's multiple comparisons test (for PDAC 95575: F=464.7, df=6, for CREP133239: F=119.9, df=6). ****p<0.0001 **e** Gating strategy for detecting effector cytokine expression. **f-g** Flow cytometric quantification of A101 CAR-T and mock-T cell expression of IFN-$\gamma$, IL-2 and TNF-$\alpha$ at E:T=50, 24hrs after coculture. n = 1. Gating strategy is shown in **e**. **h-k** Cytotoxic activity of human T cells when co-cultured with human tumor cells. The cell lines tested were A431 (**h**), A431/H9 (**I**), KLM1 MSLN KO (**j**) and KLM1 tumor cells (**k**). Data points are mean ± SE. Statistical analysis was performed using two-way ANOVA with Tukey's multiple comparisons test. For A431; n=2 independent experiments, A101 CAR-T vs mock-T; ****p<0.0001 (F=26.96, df=3). For A431/H9; n=4 (A101) and n=2 (YP218) independent experiments. A101 CAR-T vs mock-T; ****p<0.0001, YP218 CAR-T vs mock-T; ****p<0.0001 (F=83.24, df=3). For KLM1 MSLN KO; n=4 independent experiments, A101 CAR-T vs mock-T; ***p=0.0002, YP218 CAR-T vs mock-T; *p=0.0186 (F=8.851, df=3). For KLM1; n=6 independent experiments, A101 CAR-T vs mock-T; ****p<0.0001, YP218 CAR-T vs mock-T; ****p<0.0001 (F=54.99, df=3).

Supp fig 3.

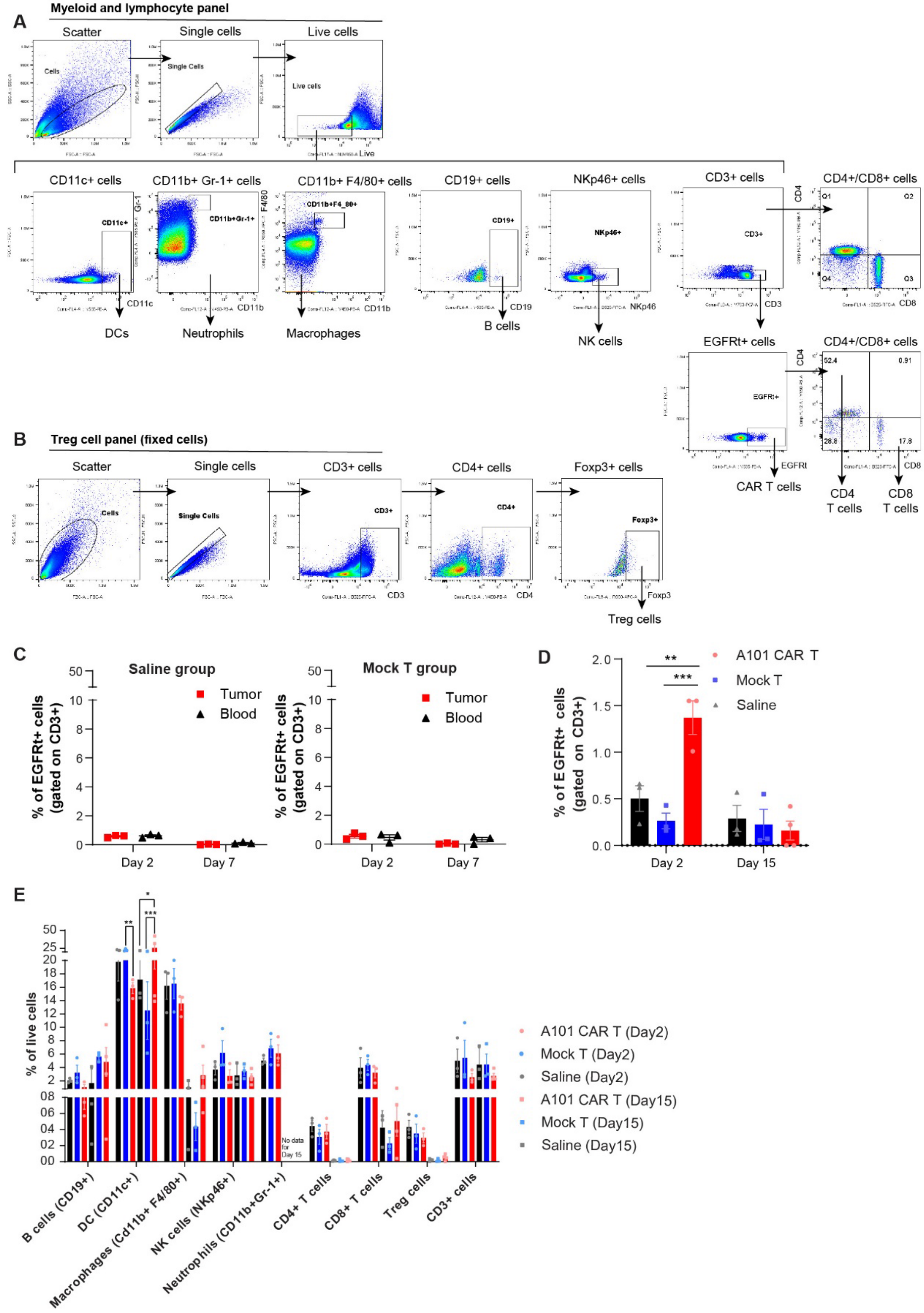

A
Myeloid and lymphocyte panel
Scatter
Single cells
Live cells
CD11c+ cells
CD11b+ Gr-1+ cells
CD11b+ F4/80+ cells
CD19+ cells
NKp46+ cells
CD3+ cells
CD4+/CD8+ cells
DCs
Neutrophils
Macrophages
B cells
NK cells
EGFRt+ cells
CAR T cells
CD4 T cells
CD8 T cells
B
Treg cell panel (fixed cells)
Scatter
Single cells
CD3+ cells
CD4+ cells
Foxp3+ cells
Treg cells
C
Saline group
Mock T group
Tumor
Blood
% of EGFRt+ cells (gated on CD3+)
Day 2
Day 7
D
A101 CAR T
Mock T
Saline
% of EGFRt+ cells (gated on CD3+)
Day 2
Day 15
E
% of live cells
No data for Day 15
A101 CAR T (Day2)
Mock T (Day2)
Saline (Day2)
A101 CAR T (Day15)
Mock T (Day15)
Saline (Day15)
B cells (CD19+)
DC (CD11c+)
Macrophages (Cd11b+ F4/80+)
NK cells (NKp46+)
Neutrophils (CD11b+Gr-1+)
CD4+ T cells
CD8+ T cells
Treg cells
CD3+ cells

**Supplementary Fig. 3. Characterization of TME changes in response to A101 CAR-T cell treatment. a-b** Gating strategy cell population analysis. **c** Percentage of $EGFRt^+$ cells among $CD3^+$ cells in tumors and blood of the control groups. **d-e** Treatment of PDAC 95575 syngeneic tumor bearing C57BL/6NCrL mice with 7 x $10^6$ A101 CAR-T cells, mock-T cells or saline. Tumors were harvested on day 2 and 15 post treatment and characterized by flow cytometry. Live cells from tumor tissue were column sorted prior to flow cytometry. **d** The percentage of $EGFRt^+$ CAR-T cells among $CD3^+$ cells. Data points are mean ± SE (for day 2: n = 3, for day 15: n = 4 for CAR-T, n = 3 for mock-T and saline) and groups were compared using two-way ANOVA analysis with Tukey's multiple comparison tests (F=8.076, df=2). ***p=0.0002, **p=0.0019. **e** Cell population analysis of PDAC 95575 tumors. Data points are mean ± SE (for day 2: n = 3, for day 15: n = 4 for CAR-T, n = 3 for mock-T and saline) and groups were compared using two-way ANOVA analysis with Tukey's multiple comparison tests (day 2: F=7.059, df=2, day 15: F=1.424, df=2). **p=0.0011, ***p=0.0001, *p=0.0171. Gating strategy for (**d**) and (**e**) is shown in **a** and **b** respectively.

Supp fig 4.

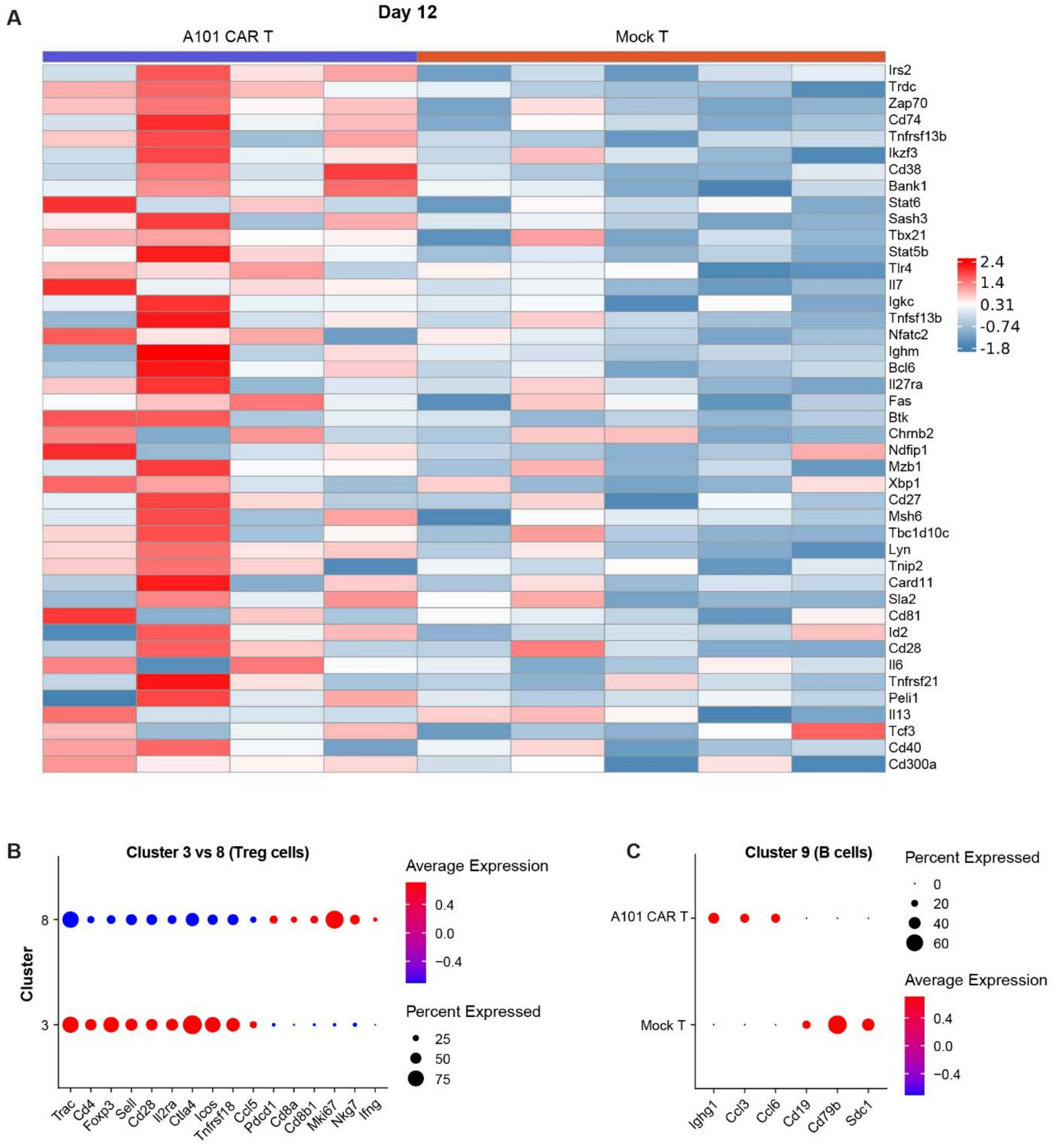


**Supplementary Fig. 4. Total and scRNA-seq data analysis of 344SQ tumors. a** Total RNA-seq data shows B cell activation on day 12 post treatment in A101 CAR-T-treated 344SQ tumors. Heat map of GSEA leading edge genes associated with the BP:GO_regulation_of_B_cell_activation pathway in A101 CAR-T- or mock-T-treated tumors on day 12 post treatment (ES=0.44, NES=1.89, pval=0.00036, padj=0.014). **b-c** Dot plots visualize gene expression in indicated clusters. Red indicates a higher average expression of the gene, and a larger dot size indicates that the gene is present in a larger percentage of cells within that cluster. **b** scRNA FLEX-seq identifies two Treg cell subpopulations in 344SQ tumors. Scaled expression of significant, differentially expressed genes in the indicated clusters ($p<0.05$). **c** Comparison of Cluster 9 cells (B cells) between CAR-T and mock-T-treated 344SQ tumors. Scaled expression of significant, differentially expressed genes ($p<0.05$).

**Supplementary Table 1. List of reagents and equipment.**

| Reagent or Resource | Source | Identifier |
|---|---|---|
| **Antibodies** | | |
| Anti-mouse CD16/32 (Clone 93) | BioLegend | Cat#: 101302; RRID: AB_312801 |
| A101 hFc | Dr. Mitchell Ho, NIH | N/A |
| PE goat anti-human IgG (polyclonal) | Jackson Research | Cat#: 109-116-170; RRID: AB_2337681 |
| FITC anti-mouse CD3 (Clone 145-2C11) | Invitrogen | Cat#: MA5-17658; RRID: AB_2539048 |
| PE anti-human EGFR (Clone Hu1) | R&D Systems | Cat#: FAB9577P; RRID: AB_2942015 |
| APC anti-mouse CD45 (Clone I3/2.3) | BioLegend | Cat#: 147708; RRID: AB_2563539 |
| iFluor 647 anti-camelid $V_HH$ (Clone 96A3F5) | GenScript | Cat#: A01994; RRID: none |
| PE human IgG1 Isotype (Clone QA16A12) | BioLegend | Cat#: 403503, RRID: AB_3097044 |
| eFluor 450 anti-mouse CD4 (Clone RM4-5) | eBioscience | Cat#: 48-0042-82, RRID: AB_1272194 |
| PE/Cyanine5 anti-mouse CD4 (Clone GK1.5) | BioLegend | Cat#: 100409; RRID: AB_312694 |
| FITC anti-mouse CD8 (Clone 53-6.7) | BioLegend | Cat#: 100706; RRID: AB_312745 |
| PE anti-mouse CD25 (Clone PC61) | BioLegend | Cat#: 102007; RRID: AB_312856 |
| APC anti-mouse Foxp3 (Clone FJK-16s) | BioLegend | Cat#: 17577382; RRID: AB_469457 |
| PE/eFluor™ 610 anti-mouse CTLA-4 (Clone UC10-4B9) | eBioscience | Cat#: 61-1522-82; RRID: AB_2574581 |
| APC anti-mouse Lag-3 (Clone eBioc9B7W) | eBioscience | Cat#: 17-2231-80; RRID: AB_2573183 |
| PE-Cyanine7 anti-mouse PD-1 (Clone J43) | eBioscience | Cat#: 25-9985-80; RRID: AB_10853672 |
| eFluor 450 anti-mouse Tim-3 (Clone 8B.2C12) | eBioscience | Cat#: 48-5871-80; RRID: AB_2574080 |

**Supplementary Table 1. List of reagents and equipment.**

| | | |
|---|---|---|
| APC anti-mouse IL-2 (Clone JES6-5H4) | Invitrogen | Cat#: 17-7021-81; RRID: AB_469489 |
| PE-Cyanine7 anti-mouse IFN-γ (Clone XMG1.2) | Invitrogen | Cat#: 25-7311-82; RRID: AB_469680 |
| eFluor 450 anti-mouse TNF-α (Clone MP6-XT22) | Invitrogen | Cat#: 48732180; RRID: AB_1548828 |
| Anti-mouse CD3 (Clone CD3-12) | Bio-Rad Laboratories | Cat#: MCA1477; RRID: AB_321245 |
| PE-Cyanine7 anti-mouse CD3 (Clone 17A2) | BioLegend | Cat#: 100219; RRID: AB_1732068 |
| VioBlue anti-mouse CD11b (Clone M1/70.15.11.5) | Miltenyi | Cat#: 130-113-238; RRID: AB_2726047 |
| PE anti-mouse CD11c (Clone N418) | Miltenyi | Cat#: 130-122-952; RRID: AB_2801981 |
| PE anti-mouse CD19 (Clone 6D5) | Miltenyi | Cat#: 130–102-598; RRID: AB_2661112 |
| APC anti-mouse F4/80 (REA126) | Miltenyi | Cat#: 130-116-525; RRID: AB_2733417 |
| FITC anti-mouse NKp46 (Clone 29A1.4.9) | Miltenyi | Cat#: 130–102-300; RRID: AB_2661345 |
| APCeF780 anti-mouse Gr-1 (Clone RB6–8C5) | Miltenyi | Cat#: 47–5931-80; RRID: AB_1518805 |
| **Chemicals, peptides, recombinant proteins and assays** | | |
| Dynabeads Mouse T-Activator CD3/CD28 for T-Cell Expansion and Activation | Gibco | Cat#: 11453D |
| Cell lysis buffer | Promega | Cat#: E1941 |
| LIVE/DEAD™ Fixable Aqua Dead Cell Stain Kit, for 405 nm excitation | Invitrogen | Cat#: L34957 |
| Foxp3/Transcription Factor Staining Buffer Set | eBioscience | Cat#: 00-5523-00 |
| 2β-mercaptoethanol | Sigma-Aldrich | Cat#: M6250 |
| DNase I recombinant, RNase-free | Roche | Cat#: 10104159001 |
| Fetal Bovine Serum | GemCell | Cat#: 100-500 |
| Penicillin-Streptomycin | Gibco | Cat#: 15140-122 |
| L-Glutamine | Gibco | Cat#: A2916801 |

**Supplementary Table 1. List of reagents and equipment.**

| | | |
|---|---|---|
| Sodium Pyruvate | Sigma-Aldrich | Cat#: S8636 |
| DMEM, high glucose, GlutaMAX™ Supplement | Gibco | Cat#: 10566016 |
| BD Cytofix/Cytoperm™ Fixation/Permeabilization Kit | BD Pharmingen | Cat#: 554714 |
| Human IL-2 | Prometheus Therapeutics & Diagnostics | Cat#: NDC 65483-116-07 |
| Mouse IFN-gamma Quantikine ELISA Kit | R&D Systems | Cat#: MIF00 |
| RNaseZap™ RNase Decontamination Solution | Invitrogen | Cat#: AM9782 |
| RNAlater™ Stabilization Solution | Invitrogen | Cat#: AM7021 |
| RBC lysis buffer | BioLegend | Cat#: 420301 |
| Non-essential Amino Acid Solution | Sigma-Aldrich | Cat#: M7145 |
| EDTA | Sigma-Aldrich | Cat#: E7889 |
| BSA | Cell Signaling | Cat#: 9998s |
| Liberase | Sigma-Aldrich | Cat#: 5401020001 |
| Lentiblast | OzBiosciences | Cat#: LBPX1500 |
| MycoAlert® Mycoplasma Detection Kit | Lonza | Cat#: LT07-318 |
| Luciferase Assay System | Promega | Cat#: E1501 |
| Pan T cell isolation kit II, mouse | Miltenyi | Cat#: 130-095-130 |
| Leica Bond Polymer Refine Kit | Leica Biosystems | Cat#: DS9800 |
| OneComp eBeads™ Compensation Beads | ThermoFisher Scientific | Cat#: 01-1111-42 |
| CD45 (TIL) MicroBeads, mouse | Miltenyi | Cat#: 130-110-618 |
| Dead cell removal kit | Miltenyi | Cat#: 130-090-101 |
| Chromium Next GEM Single Cell Fixed RNA sample preparation kit | 10x Genomics | Cat#: 1000414 |
| RNeasy Mini Kit | Qiagen | Cat#: 74104 |
| ArC™ Amine Reactive Compensation Bead Kit | Invitrogen | Cat#: A10346 |
| DAPI | Invitrogen | Cat#: D1306 |
| Zombie UV™ Fixable Viability Kit | BioLegend | Cat#: 423107 |
| LS columns | Miltenyi | Cat#: 130-042-401 |
| MS columns | Miltenyi | Cat#: 130-042-201 |
| **Cell lines & mouse strains** | | |
| Mouse: PDAC 95575 | Dr. Serguei Kozlov, CAPR, NIH | N/A |
| Mouse: PDAC 95575 GFP+/Luc+ | Dr. Mitchell Ho, NIH | N/A |
| Mouse: 344SQ | Jonathan M. Kurie, MD Anderson Cancer Center | N/A |

**Supplementary Table 1. List of reagents and equipment.**

| | | |
|---|---|---|
| Mouse: AB12 | Sigma-Aldrich | Cat#: 10092306-1VL |
| Mouse: AE17 | Sigma-Aldrich | Cat#:10092310-1VL |
| Mouse: Panc02 | Cytion | Cat#: 300501 |
| Mouse: 531LN2 | Jonathan M. Kurie, MD Anderson Cancer Center | N/A |
| Mouse: TC-1 | Prof. TC.Wu, Department of Pathology, SOM, JHU (NCI MTA. #49891-22) | N/A |
| Mouse: CREP133239 GFP+/Luc+ | Dr. Mitchell Ho, NIH | N/A |
| Mouse: LLC (LL/2 (LLC1)) | ATCC | Cat#: CRL-1642 |
| Human A431 | Ira Pastan, NCI | N/A |
| Human: A431/H9 | Ira Pastan, NCI | N/A |
| Human: KLM1 | Christine Alewine, NCI | N/A |
| Human: KLM1 MSLN KO | Christine Alewine, NCI | N/A |
| Mouse: 129S2/SvPasCrl | Charles River | Strain# 476 |
| Mouse: C57BL/6NCrL | Charles River | Strain# 027 |
| Mouse: B6.SJL-PtprcaPepcb/BoyCrCrl | Charles River | Strain# 564 |
| **Recombinant DNA** | | |
| GFP/Luciferase transduction: R980-M03-663 mPol2 ffLuc2-eGFP (pFUGW) | NCI, Frederick, NIH | N/A |
| Human A101 CAR plasmid | Dr. Mitchell Ho, NIH | N/A |
| Mouse A101 CAR plasmid | Dr. Chaido Stathopoulou, NIH | N/A |
| **Software and algorithms** | | |
| GraphPad PRISM 9 | GraphPad | RRID:SCR_002798 |
| FlowJo (version 10.7.2) | FlowJo LLC. | RRID:SCR_008520 |
| CytoFLEX flow cytometer | Beckman Coulter, USA | RRID:SCR_025068 |
| BioRender | BioRender | RRID:SCR_018361 |
| Halo | Indica Labs | RRID:SCR_018350 |
| Aperio ImageScope | Leica Biosystems | RRID:SCR_020993 |
| Seurat (version 4.4.0) | Seurat | RRID:SCR_007322 |
| R | R | RRID:SCR_001905 |
| STAR | STAR | RRID:SCR_004463 |
| RSEM | RSEM | RRID:SCR_000262 |
| LIMMA | LIMMA | RRID:SCR_010943 |

**Supplementary Table 1. List of reagents and equipment.**

| | | |
|---|---|---|
| CIBERSORT | Stanford University; Stanford; California | RRID:SCR_016955 |
